# Directed Evolution of Enzymes based on *in vitro* Programmable Self-Replication

**DOI:** 10.1101/2021.04.22.440993

**Authors:** Adèle Dramé-Maigné, Rocío Espada, Giselle McCallum, Rémi Sieskind, Yannick Rondelez

**Affiliations:** Gulliver UMR CNRS 7083, ESPCI Paris, Université PSL, 75005 Paris, France

## Abstract

High-throughput directed evolution, implemented in well-controlled in vitro conditions, provides a powerful route for enzyme engineering. Most existing technologies are based on activity screening and require the sequential observation and sorting of each individual variant. By contrast, approaches based on autonomous feedback loops, linking phenotype to genotype replication, enable autonomous selection without screening. However, these approaches are only possible *in vivo*, or applicable to very specific activities, such as polymerases or ligases. Here, we leverage synthetic molecular networks to create a programmable in vitro feedback loop linking a target enzymatic activity to gene amplification. After encapsulation and lysis of up to 10^7^ transformed variants, the genes present in each droplet are amplified according to the activity of the encoded enzyme, resulting in the autonomous enrichment of interesting sequences. Applied to a nicking enzyme with thermal or kinetic selection pressures, this method reveals detailed mutational landscapes and provides improved variants.

## Introduction

The growing number of technological applications of enzymes^1^, in medicine, chemistry^2^ or even information technology^3^, often requires their optimization for use in unnatural functions or conditions. Although techniques for rational engineering of proteins are improving rapidly, the explosive nature of the polypeptide sequence space, together with the intricate electronic effects that underlie catalytic efficiency, call for combinatorial methods to navigate their mutational landscapes^4–6^. High throughput *in vitro* screening methods, for example using emulsions and optical sorters ^7–11^, are appealing because they allow screening of many variants for improved function in non-biological conditions. However, these methods require relatively complex or costly sorting platforms, and their reliance on one-by-one observation and manipulation of variants still yields an unfavorable linear dependency between experimental effort and screening capacity.

In contrast to these supervised approaches, natural evolution is a batch process based on a direct positive feedback that internally connects phenotype and genotype propagation, offering much higher potential in terms of throughput. Artificial *in vitro* variants of natural evolution have been developed for the selection of ligands, and can explore very large nucleic or peptidic libraries with limited effort, because all variants are tested in parallel^12–16^. However, extending *in vitro* screen-less methods beyond simple binding affinity, in particular to selection for catalytic proficiency, remains a formidable challenge^17^. Among pioneering works on artificial selection^18^, the Compartmentalized Self-Replication (CSR) ^19^ relies on the microencapsulation of single bacteria expressing DNA polymerase variants together with gene-specific primers, followed by *in vitro* amplification of their own genes. In these conditions, only active polymerases efficiently replicate their own genes, and thus gradually come to dominate the population. This method takes advantage of the expression machinery of the bacteria, ensuring a genotype-phenotype linkage, while maintaining a screening-free *in vitro* selection step. Recently, Ellefson *et al.*^20^, introduced *ad hoc* genetic circuits into this process, widening the range of applications of the method, previously restricted to polymerases^21,22^. In Compartmentalized Partnered Replication (CPR), the target enzyme activity is artificially linked to the expression level of Taq DNA polymerase, which in turn controls its gene’s replication after lysis and PCR in droplets. The use of genetic circuits offers increased versatility and programmability of the selection process. However, in this case, the evolutionary pressure is applied inside living cells, limiting both the freedom to alter the selection conditions, and the range of activities that can be targeted^23^.

In this work, we leverage recent progress in molecular programming^24–27^ to construct an *in vitro* selection circuit to enrich enzyme libraries for specific variants. In this high-throughput approach, a transformed bacterial library is encapsulated and immediately lysed, releasing enzymatic variants and their encoding gene into individual droplets. Within each droplet, a molecular reaction network then non-linearly links the targeted enzymatic activity to the generation of small DNA oligonucleotides. These strands are used as primers in a subsequent droplet-PCR, thereby connecting enzymatic activity to the fitness of the encoding gene (Figure 1). We refer to this screening-free, *in vitro* approach as the Programmable External Networkbased Compartmentalized Self-Replication (PEN CSR). We demonstrate the approach for the selection of improved variants of Nt.BstNBI (NBI), a nicking enzyme from *Bacillus Stearothermophillus*, and a model enzyme among nickases^28^. We first show that, implemented in microfluidic droplets, the PEN CSR provides a selection efficiency close to the theoretical optimum^29^. When applied to a library of 5.10^5^ random NBI variants, a single round of PEN CSR provided a detailed map of its mutational landscapes under various pressures (neutral, kinetic or thermal). As a validation of the method, we used these fitness maps to construct a number of improved NBI variants, showing higher catalytic rate or thermal stability.

**Figure 1.**
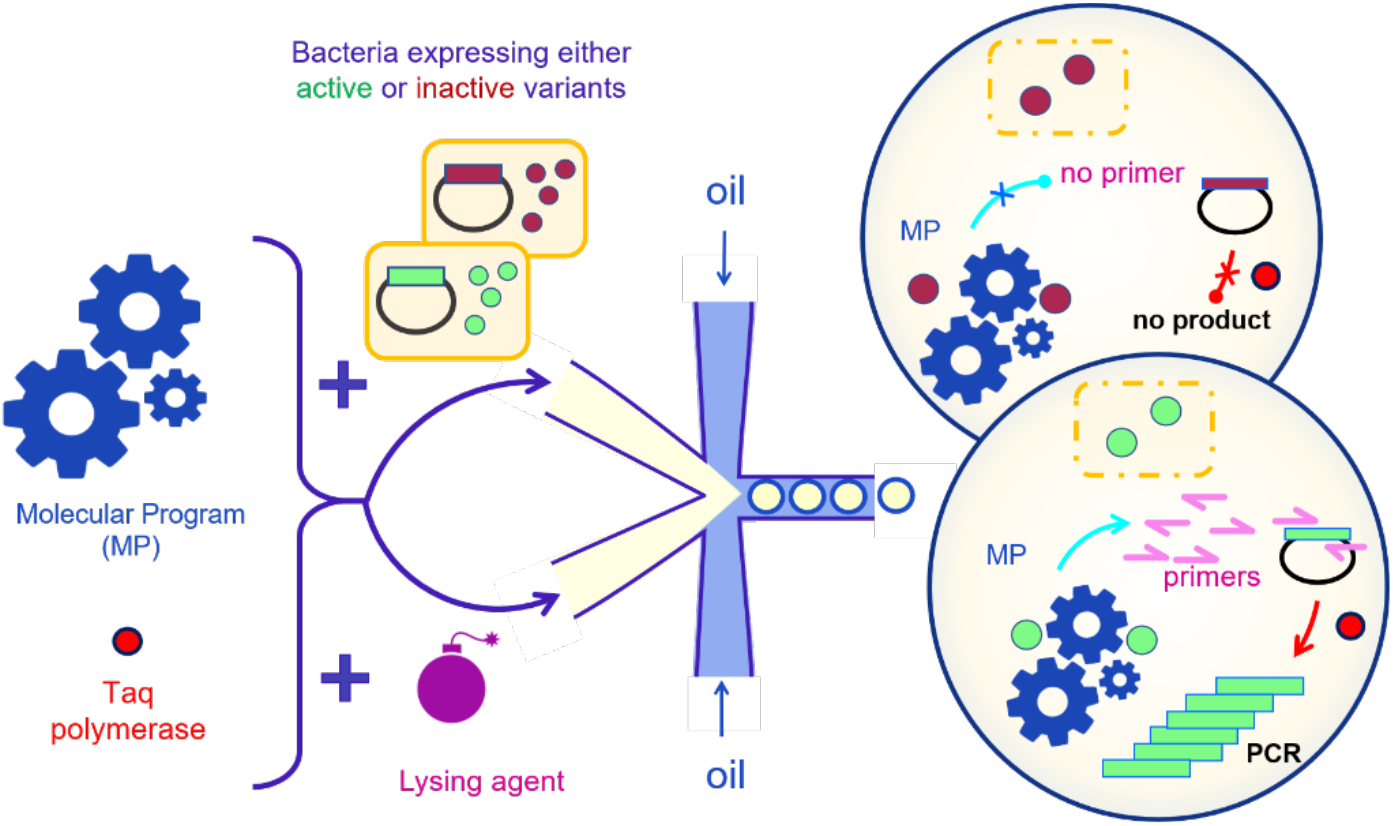
PEN CSR concept. Bacteria are transformed with a library of plasmids and express the target enzyme in vivo. The cells are compartmentalized in monodisperse water-in-oil droplets, together with a molecular program (MP), a DNA polymerase, and a lysing agent. After lysis, the molecular program autonomously evaluates *in vitro* the target enzyme activity level and accordingly generates primers against the encoding gene. The emulsion is then submitted to PCR cycles, so that only active gene variants, having triggered the generation of sufficient PCR primers, will be amplified. Overall, the fraction of active variants increases in the pooled gene population.

## Results

### Linking enzyme activity to primer production

We selected NBI as a target because of its many applications, including molecular diagnostics^30,31^ and molecular technologies^2,33^. NBI is currently used in its natural form, although many endonucleases have been engineered for better performance *in vitro*^34^. To build the autonomous selection circuit, we used the PEN DNA Toolbox, a molecular programming language based on dynamically redirecting the activity of a DNA polymerase between short templates using endogenous oligonucleotides^35,36^. Here, the goal was to connect the sitespecific NBI nickase activity to the generation of *nbi*-targeting PCR primers.

The molecular network thus performs a number of tasks: it first senses NBI activity, then amplifies and reshapes the molecular signal, and finally converts it to primer production yield (Fig. 1). Based on numerical simulations and preliminary testing, we selected a molecular circuit containing 3 stages (Fig. 2 and S1), all encoded by synthetic DNA templates. The process initiates on the *source* template (*sceT*), a hairpin loop that primes itself and displays the NBI recognition site. Upon polymerase-nickase cycles, it produces a signal oligonucleotide (*s*) linearly over time. The *non-linear* module is an autocatalytic template (*aT*) that has the same sequence as input and output and therefore encodes a positive feedback loop, which exponentially amplifies the incoming signal *s*. This quickly generates the amount of strands necessary to load the third stage, the *primer-generating* module. The primer-generating module contains two primer-producing templates *ppT_F_* and *ppT_R_*, which generate the forward and reverse primers, respectively, using the signal *s* as input. These two modules also depend on NBI activity. Altogether, after assembly of the full reaction network, we observed that this circuit (called IPA for Isothermal Primer Amplification), running at a constant temperature of 45°C, can translate the presence of a specific nickase activity into the production of a graded concentration of primers, reaching 6 nM/min at high nickase concentrations (Fig. 3A).

**Figure 2.**
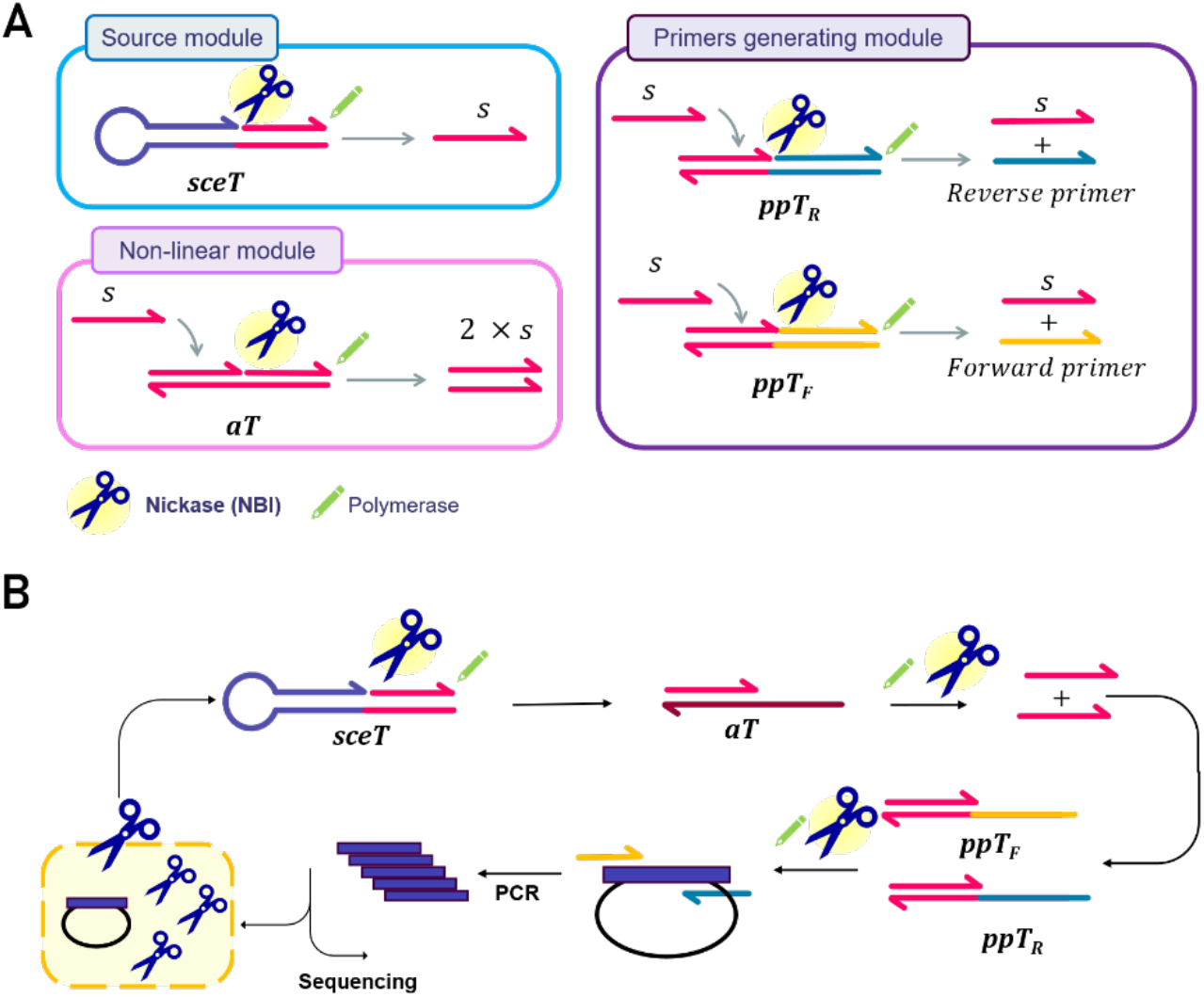
PEN CSR modules. **A**. Three modules compose the molecular circuit translating NBI activity into primers concentration. The source module generates trigger strands (*s*) from the source template (sceT). The non-linear module catalyzes the amplification of the trigger *s*. The primer-generating module receives the trigger *s* as input and produces reverse and forward primers, targeting the *nbi* gene in its expression plasmid. All modules use a thermophilic DNA polymerase and depend on specific NBI nicking activity. **B**. Module connections enabling the translation of NBI activity into PCR amplification of *nbi* gene.

**Figure 3.**
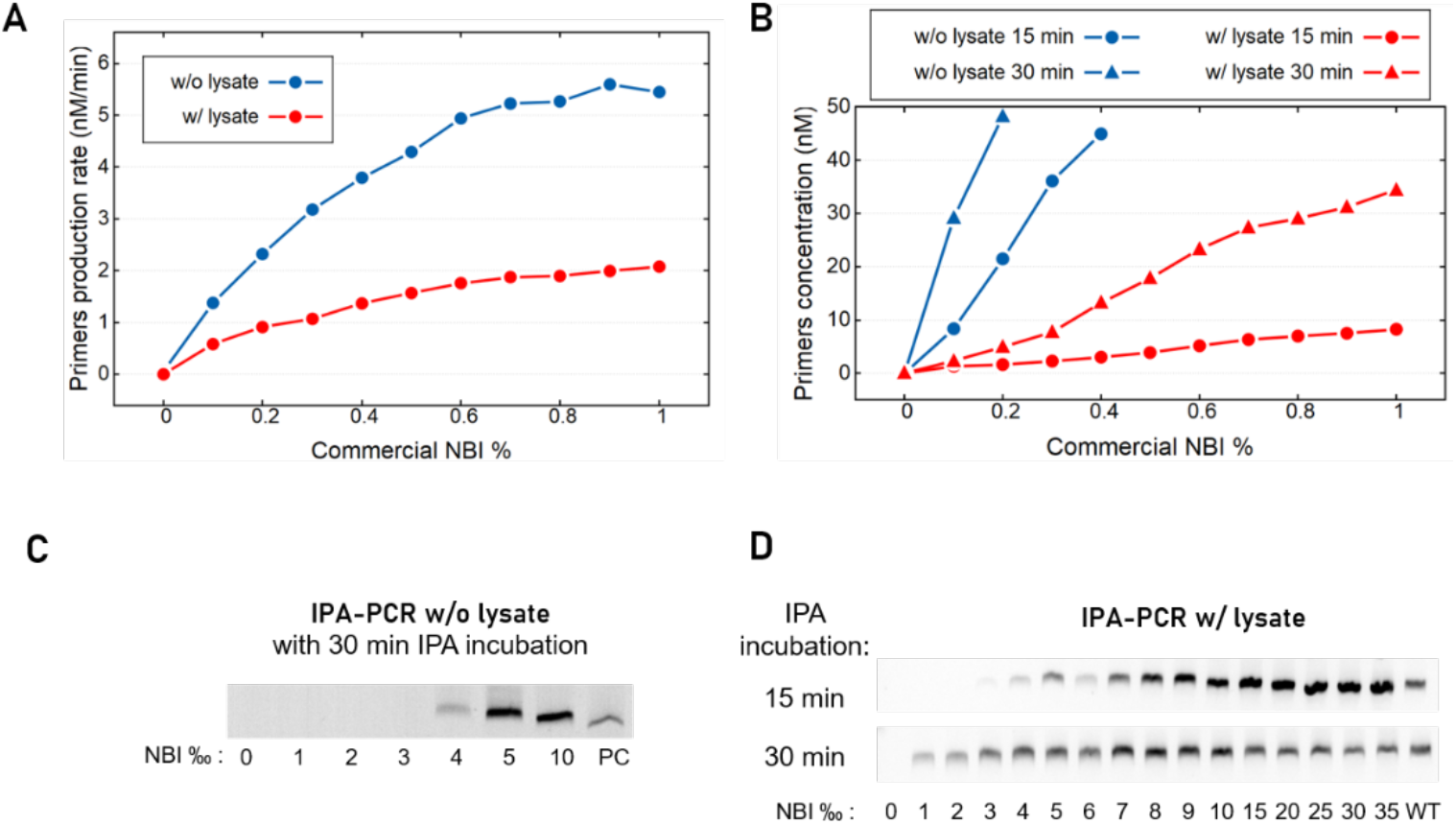
Linking enzymatic nicking activity to PCR amplification. **A**. Initial primer production rate with or without bacterial lysate at a concentration of 100 bact/nL as a function of commercial NBI concentration. **B**. Primer concentration reached after 15 or 30 min of IPA incubation with or without bacteria lysate as a function of commercial NBI concentration. **C**. Agarose gel electrophoresis of the IPA-PCR product after 30 min IPA incubation with various commercial NBI concentrations and without bacteria lysate. PC indicates an experiment with 500 nM of spiked-in primers **D**. IPA-PCR product after 15 or 30 min IPA incubation with bacteria lysate (100 bact/nL). WT indicates lysate from bacteria expressing the wild type enzyme.

### Linking enzyme activity to PCR amplification

To connect the IPA activity-to-primers converter to a *nbi* gene amplification process, the reaction mixture was complemented with a PCR polymerase and submitted to thermal cycles following isothermal incubation. Because all stages of the selection process must ultimately be performed in the same closed droplet, we optimized reaction conditions and primer design (melting temperature around 58°C and ~20 bases length, Fig. S2-S3), such that both the IPA and PCR steps could be run in the same buffer. Primer binding sites were selected ~150 bases upstream and downstream of the gene, to allow a nested recovery of the genetic information following droplet PCR. A robust NBI-dependent *nbi* gene amplification was observed in these conditions (Fig. 3C).

When attempting to perform this reaction in the presence of bacterial lysate, as would be the case in the droplet setup, we found that both the IPA and the PCR steps were strongly affected by components released from lysed cells, even at bacteria concentration as low as 50 bact/nL (~1:20000 dilution factor). We therefore protected the molecular program templates from exonuclease degradation using chemical modifications (phosphorothioate bonds^35^ and terminal biotinylation in conjugation with streptavidin (Fig. S4) and selected a lysate-resistant PCR polymerase (OneTaq polymerase, Fig. S5). Primer production rate decreased in the presence of lysate (Figure 3A and B), but was sufficient to perform the coupled IPA-PCR reaction (Fig 3D). This reaction was possible with up to ~150 bact/nL (Fig. S6), corresponding to 1.5 bacteria per 10 pL, our target droplet volume. PCR yield was dependent on both nickase activity and IPA incubation time (Fig. 3D), suggesting the possibility to tune the stringency of our selection module through the time of incubation.

### Interfacing with bacterial gene expression

To complete the self-selection loop, we used a T7 express® bacterial strain to express the library of NBI nickases from their encoding genes, prior to droplets encapsulation and IPAPCR. To avoid nickase toxicity towards the bacterial genome in the expression strain, we relied on the leaky expression of a cognate methylase (Ple1) from a high copy plasmid, as generally described for restriction enzyme expression^37^. We optimized the concentration of an enzymatic lysing agent (rLysozyme) in our reaction mixture to lyse cells in a controlled and droplet-compatible manner following induction of nickase expression (Fig. S6). Thus, the expressionlysis-IPA-PCR chain provided a complete one-pot genotype-to-phenotype-to-genotype feedback loop, applicable to the autonomous selection of nicking activities in droplets. We tested primer production from NBI-expressing bacterial lysate at relevant dilutions, both for the wild-type NBI and the inactive variant E418A^28^ (Fig. 4A). High values were obtained only for the active variant, with bacterial concentration between 50 and 100 bact/nL. We then ran the full process in bulk and observed an NBI activity-dependent amplification of *nbi* gene for various IPA incubation times, ranging from 15 to 60 min, while no amplification was observed for the inactive NBI (Fig. 4B).

**Figure 4.**
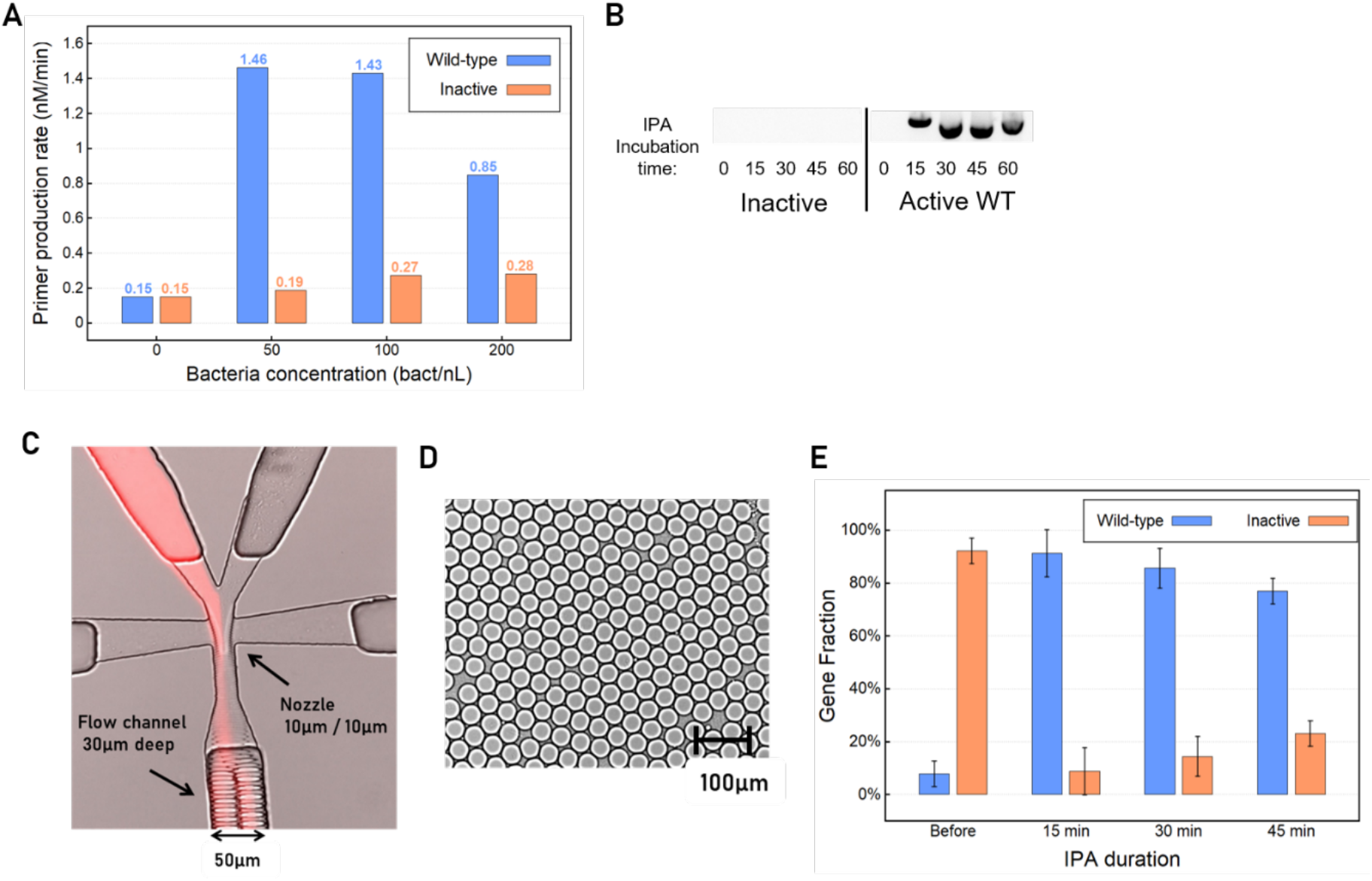
Autonomous selection of a mock bacterial library. **A**. Primer production rate measured in lysates from bacteria expressing either the wild-type, or an inactivated point variant, according to bacteria concentration. **B**. Agarose gel electrophoresis of the Lysis-IPA-PCR process after 0 to 60 min of IPA incubation at 100 bact/nL, for bacteria expressing either the active wild-type or the inactive variant. **C**.Flow-focusing junction. The bacteria are injected from the right aqueous inlet, while the lysing agent is present in the left aqueous inlet, together with a red-fluorescent dye used for flow-balancing purposes. **D**.The microfluidic device generates droplets of ~26 μm. **E**. Quantification of gene frequency upon one round of autonomous selection, starting from a 1:10 mixture of wild-type and inactivated variant.

### Quantifying selection efficiency

To test the efficiency of this method to perform compartmentalized self-selection, we created a mock bacterial library containing a fraction of active *nbi* genes diluted in a majority of inactive variants. We designed a 2-layer^38^ microfluidic flow-focusing device, with two inlets, enabling separate injection of cells on one side, and lysing agent on the other, thus avoiding release of enzymes before encapsulation (Fig. 4C and S7). 10^7^ monodisperse droplets (27.6 +/- 0.8 μm diameter, volume 9.9 pL, Fig. 4D) were produced at the oil junction in ~30 minutes (rate 6.6 kHz) and a mean occupancy of ~0.4 cell per droplet. The droplet population was collected at the outlet, submitted to the IPA step at 45°C for 15, 30 or 45 min, followed by the PCR thermal protocol. The emulsion was then broken and the water phase was collected. We used an rhqPCR assay^39^ to measure the fraction of active and inactive genes before and after selection. Whereas the initial library contained only 10% of active clones, they were enriched to ~80-90% in the product of *in vitro* autonomous selection, with shorter IPA incubation time accounting for higher final active gene fraction (Fig. 4E). We used these measurements to compute the Selection Quality Index (SQI)^29^, a metric that considers the poissonian distribution of bacteria in the droplets to evaluate the selection efficiency. With SQIs between 0.9 and 1, this corresponds to an almost optimal selection protocol.

### Selection for improved kinetics and thermostability

We applied this approach to scan the entire 1.8 kb *nbi* wildtype gene (604 amino acids), in search of positions harboring interesting mutations. In addition to a neutral selection experiment (30 min IPA), we designed and implemented two different selection pressures, reflecting two desirable improvements of the NBI enzyme with respect to its applications: increased kinetic rate (Kinetic Stress, KS) and increased thermostability (Thermal Stress, TS). To do so, we first modified the selection protocol in such a way that the wild-type enzyme would *not* produce a detectable amount of PCR product under the selective conditions (Fig. S8). For the KS conditions, we decreased the IPA incubation time to 10 minutes. For the TS selection, we applied a 3 min heat shock at 65°C, a previously observed critical temperature^40^, before the usual 30 min IPA incubation. We created a randomized library using error-prone PCR at ~3.5 mutations per kb and obtained a cloned library with a size of ~5.10^5^, showing 20% of the wildtype activity, as measured using a fluorescent hairpin reporter oligo (Fig. S9). This library was submitted to selection in the neutral, KS or TS conditions, with a coverage of 20x (approximately 10^7^ cells tested) in each case, in less than 3 hours. Subsequent qPCR quantification showed that DNA amplification fold was higher in droplets than in controls in which the reaction was performed without micro-compartmentalization. Following selection and re-cloning, the enriched libraries displayed 2x and 1.7x increase in activity compared to the initial library in the KS and TS conditions, respectively, indicating that the randomized library had been successfully submitted to purifying and/or positive selection in both cases (Fig. S10).

### Pressure-specific favorable mutations revealed by mutational scanning

The initial and final libraries were sequenced using a nanopore MinION device, obtaining more than 10^5^ full length reads for each set (Fig. S11). After data polishing using 23095 wild-type reads as reference^41^, we detected negative and positive selection as a negative or positive shift in averaged mutation frequency between each final library and the initial one. We computed a fitness effect F_eff_ for each mutation as F_eff_ = Ln(f_f_/f_i_), where f_i_ and f_f_ are the initial and final frequency for each mutation, averaged over the polyclonal population^42^. This provided mutational F_eff_ estimations ranging from −1.3 to +2.3, with reasonable bootstrapped uncertainty estimate (95% confidence interval < 0.5) for more than 3200 mutations in each case (i.e. 28% of all 604*19 possible mutations, Fig. 5A,B and S12-14). Among these, 137 and 38 nonsynonymous amino acid replacements had clear positive effects (p-value <.05) in the KS and TS selections, respectively (Fig. S15). Computing the relative entropy for each position as a measure of local mutational sensitivity^7^ provided clear structural patterns, where sensitive positions clustered around the DNA binding pocket for the neutral and KS selections, but were more distributed over exposed residues for TS. This analysis also clarifies previous attributions of the active site residues (Fig. 5C,D, S16-17 and SI movies 1-3).

**Figure 5.**
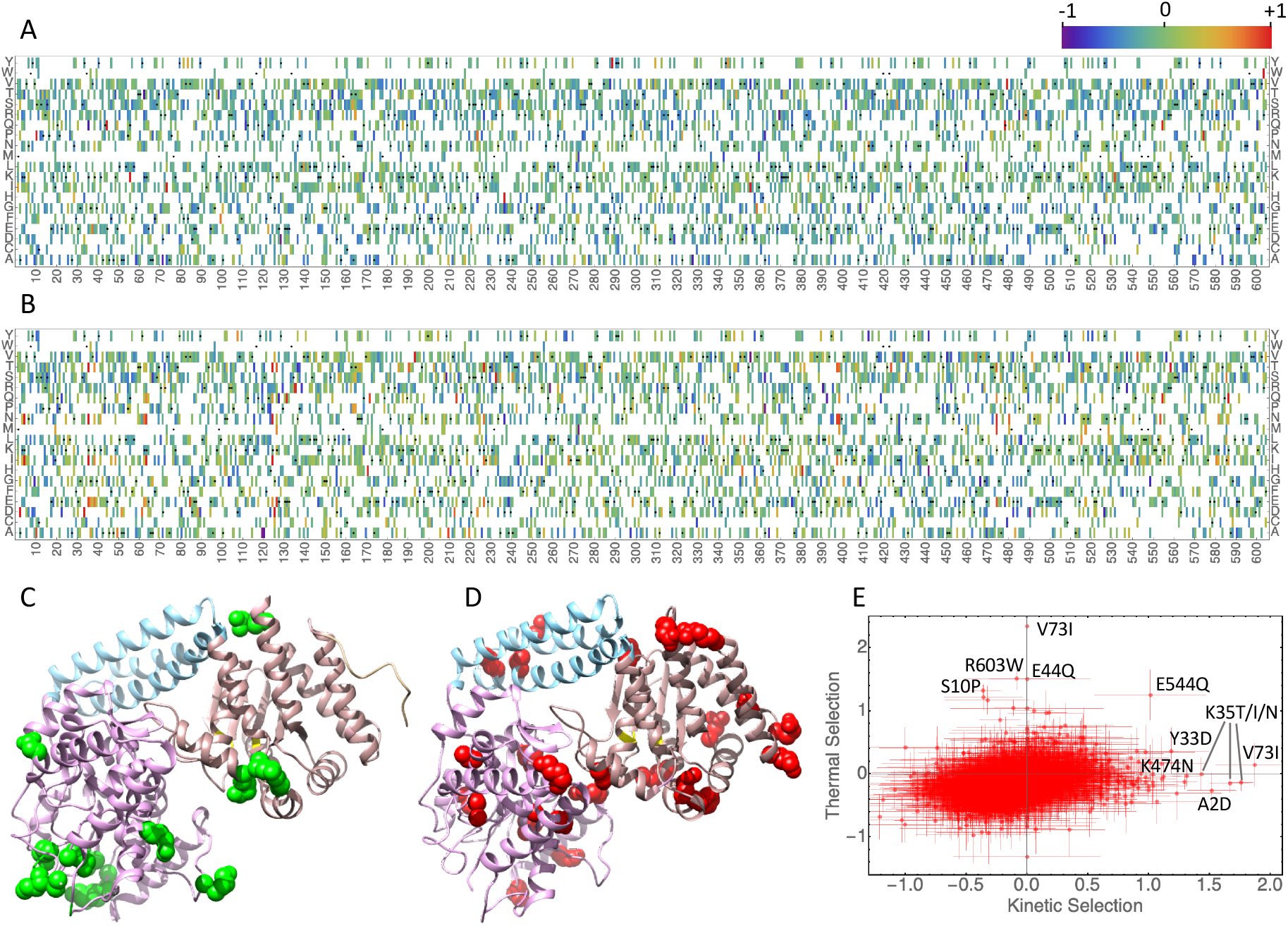
Mutational scanning of NBI under selective pressure. **A**, **B**. Computed fitness effect F_eff_ for 28% of possible single mutations in TS (**A**) and KS (**B**). Black dots indicate the wild-type amino acid at a particular position. **C**, **D** The 25 most favorable mutations in the kinetic selection (green, KS) or thermal selection (red, TS) are shown on the structural model of NBI (based on PDB 2ewf). In pink the recognition domain, in blue the linker domain and in light brown the catalytic domain with putative catalytic amino acids in yellow. **E**. TS versus KS F_eff_ scores.

The 25 most favorable mutations in the kinetic selection (A2D, K35N/I/T/E, R558T, K4N/I, G125D, R232S, K124I/T/N, K126E, Y33D, K63N, K397N, K474M/N/E, K398N, D170H, K63I, V84E, G212D) are shown in green on the structural model of the enzyme in Fig. 5C. It is striking that among these, 19 (76%) affect lysine or arginine residues lining the DNA binding interface^43^. This suggests that the dominant mechanism behind the nickase’s increased turnover is a decreased electrostatic interaction with DNA. This observation is consistent with previous observation of product inhibition affecting nicking enzymes^44^. However, the remaining mutations affect different parts of the enzyme, suggesting that other mechanisms exist which can also increase the turnover rate. For TS conditions, in contrast, the strongest effects were distributed throughout the structure (V73I R603W S10P E44Q K187H Q55K T574P H288Y Q478Q L355I E554Q N531K F479V L99F V37E D236H N412N R488R L432I N344Y A163S K581I E579D V139L V358G, Fig. 5D), were less biased in terms of amino acid chemistry or protein domain, and did not tend to aggregate on hotspot positions (0 versus 12 mutations were colocalized with at least another in TS and KS, respectively). Interestingly, three of these 25 beneficial mutations are synonymous replacements, suggesting that increased expression may also be selected by the TS protocol. We checked the relative distribution of synonymous mutation effects (Fig. S18) and found that codon usage appeared indeed more constrained in the TS protocol. Additionally, beneficial synonymous replacements in TS do not translate in KS. These observations, consistent with the plateauing of the PCR yield at higher wild-type NBI concentration (Fig. 3D), suggest that KS selects for true increases in specific activity, with limited bias brought by higher protein expression.

More generally, we found that beneficial mutations in the two sets are almost orthogonal, with most of the best mutations for one property being close to neutral on the other (Fig. 5E). Some exceptions exist, such as E554Q, which appears beneficial in both cases, although it is located at the beginning of an alpha helix of the catalytic domain, far away from the active site and the DNA binding site.

### Confirmation of mutation effects

To validate these results, we expressed and quantified the activity of several NBI variants containing beneficial mutations from the TS and KS selections. From the mutations enriched in the KS protocol, we expressed the single mutants A2D, K35N, K474N, R558T, and the double mutants K124N/K474N, K35N/R558T. From the mutations enriched in the TS protocol, we expressed the single mutants E44Q, V73I, R603W, the double mutant E44Q/R603W, and the triple mutant S10P/E44Q/R603W. All expressed variants showed activity in cell lysate (Fig. 6A,B). Most variants with mutations from the KS set were faster than the wild-type enzyme o a test substrate (Fig. S9), with double mutants K35N/R558T and K124N/K474N lysates being twice and three times more active, respectively (Fig. 6A). We evaluated the thermal resistance of all variants by incubating the samples at 65°C for 3 minutes before measuring their activity. Those with KS mutations lost most of their activity, while TS variants had a thermal resistance similar to or higher than the wild-type enzyme (Fig. 6B). In some cases, such as R603W, the gain in remaining activity originates mostly from a higher initial value in the lysate. This could indicate an improvement in the expression level of this mutant, rather than a change in its enzymatic property. By contrast, the double and triple mutants E44Q/R603W and S10P/E44Q/R603W have an initial activity similar to the wild type but are less affected by the heat shock. The triple mutant in particular shows almost no loss during incubation (activity ratio~ 82%).

**Figure 6.**
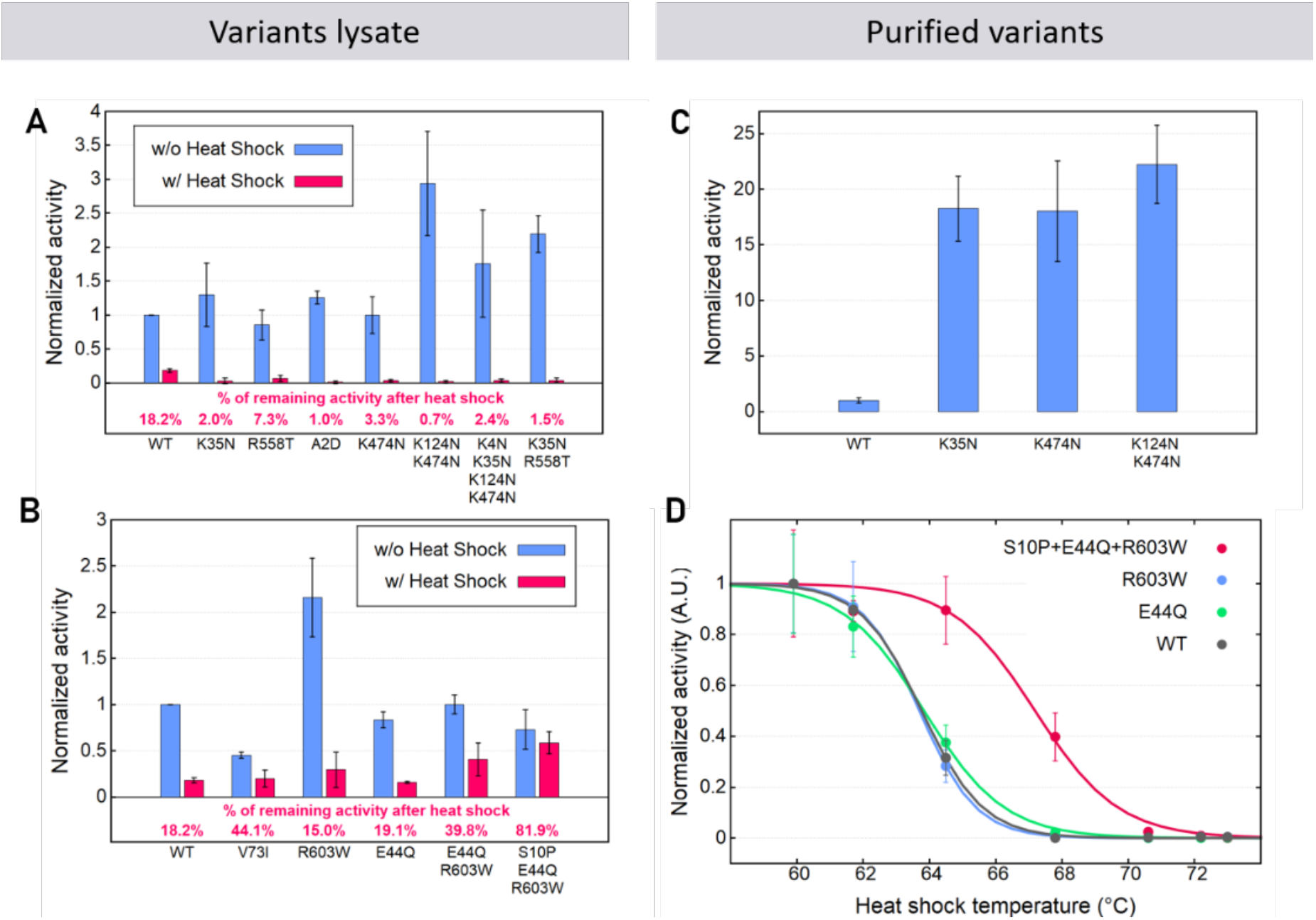
Activity measurements of reconstructed NBI mutants. **A, B** Measurements in cell lysate, at constant concentration of bacteria (25 bact/nl). Activity normalized with respect to the wild-type (WT) activity of variants containing mutations arisen in the KS (**A**) or the TS (**B**) experiment with (w/) or without (w/o) 3 min heat shock at 65°C. **C**, **D**, Measurements for some purified protein variants, at constant protein concentration. **C**. Activity normalized with respect to the wild-type (WT) activity of purified variants containing beneficial mutations from the KS experiment. **D**. Remaining activity ratio after 3 min incubation for variants containing mutations selected by the TS protocol. The data points are fitted with a sigmoid.

To disentangle enzymatic properties from their protein expression levels, we purified representative variants and measured their activity at normalized concentrations. In these purified conditions, the 3 variants from the KS set (K35N, K474N, K124N/K474N) showed a significant increase in their activity, between 18 and 22-fold (Fig. 6C), while retaining at least partial specificity (Fig. S19). We constructed thermal stress curves for E44Q, R603W and S10P/E44Q/R603W (Fig. 6D), in the presence of nonspecific DNA (Fig. S20). T_1/2_ for R603W and E44Q (63.7 °C and 63.8 °C respectively) are similar to the wild type (T_1/2_ = 63.8 °C). This is consistent with the hypothesis that the KS protocol also favors higher expression. In the case of the triple mutant, however, we observed a substantial thermal stabilization (T_1/2_= 67.2 °C).

## Discussion

Molecular programming is an emerging field which leverages synthetic informational polymers to encode complex behaviors *in vitro*. Here, we demonstrate a new application for this field, where a synthetic circuit is created to apply a pre-programmed selective pressure on a catalytic target. In combination with microencapsulation, we generate an all-in-one *in vitro* directed evolution platform, acting as an autonomous filter to select catalytic variants in an enzymatic library. The use of a synthetic molecular program not only offers the possibility to select in non-physiological conditions (exemplified here with the thermal stress selection), but also increases the degrees of freedom in the design of the selection scheme. For example, it has been suggested that the shape of the selection function, which we can adapt by tuning the IPA step (Fig. 3 and S1), is a key driver of the efficiency of the evolutionary exploration of a fitness landscape^45^. Additionally, more complex selection networks could be built to perform smarter selection operations, for example integrating logic or neural operations^46^, in order to take several enzymatic features into account for the scoring of a given variant. An appealing application would be to directly evolve enzymes with tailored selectivity, that is, active on one reaction AND inactive on others, a process that requires alternate selection and counterselection cycles with classical methods.

Our strategy demonstrates versatile and high throughput *in vitro* evolution, while avoiding any advanced microfluidic operation such as sorting, fusion, reinjections, etc.^47^ A corollary is that lysis, IPA and PCR have to work as a one-pot multi-step process in a single droplet, which requires fine tuning of the reaction conditions. In particular, we found that the cell lysate fragilized the subsequent *in vitro* steps (IPA and PCR), which had to be specifically optimized. However, it is likely that the mitigating strategies introduced here can be reused for other evolution campaigns using PEN CSR. Notably, the limited range of bacteria concentration in which our system works creates a built-in mechanism that enhances the efficiency of the selection process: because the IPA-PCR reaction cannot run in droplets containing two or more bacteria, the carry-over of inactive variants via co-encapsulation is limited^29^, likely contributing to the exceptional performance of the selection process.

We believe that programmable self-selection implemented with PEN CSR will greatly enlarge the variety of enzymes that can be evolved *in vitro* using a screening-free strategy. While it appears generally straightforward to couple most DNA-processing enzymes to a DNA based molecular program^48^, we note that natural^49^ or synthetic^50^ strategies also exist to connect the presence of various small molecules, and thus the corresponding enzymes, to nucleic regulatory processes and circuits.

In principle, self-selection networks open the route to extremely high throughput evolution, since they remove the need for individual evaluation of every mutant in a library. Here, we found the throughput to be limited by bacterial transformation efficiency using cloned DNA. By coupling the PEN CSR technology and an *in vitro* enzyme expression platform^51^, a wide range of toxic proteins could be targeted, while also avoiding the transformation bottleneck.

Finally, in combination with Next-Generation Sequencing, PEN-CSR provides an efficient way to explore the mutational landscape of enzymes. Our results extend the observation, previously made on ligand evolution^52^, that a single round at sufficient throughput is sufficient to reveal beneficial mutations, although each of them are represented typically at a frequency of less than 0.1% in the selected libraries (Fig. S15). The analysis we made here was based on the simple assumption of independence between mutational effects^42^. Since we designed our approach to recover full length sequences, we anticipate that its coupling with less naïve, epistatic models^53^ will allow a better exploitation of the data it generates.

## Acknowledgements

This research was funded by the European Research Council (ERC-CoG-2014 ProFF ID: 647275), PhD fellowships from French Ministry of Higher Education and Research (CDSN ENS Lyon and Paris-Saclay resp.) to A.D.-M. and R.S. and postdoctoral fellowship from European Union’s Horizon 2020 research and innovation program (Marie Skłodowska-Curie No 845976) to R.E. The microfluidics was realized at the Pierre-Gilles de Gennes Institute (IPGG) with the support of the Equipex/Idex ANR program (Equipex ANR: 10-EQPX-0034). We thank T. Fujii, his laboratory and the Institute of Industrial Science (IIS) of the University of Tokyo for hosting the first steps of the project, V. Taly for the sample of EA-surfactant, R. Hsu and A. Kazuaki for preliminary work, D. Kiga and G. Gines for advices and several colleagues for fruitful discussions and helpful comments on the manuscript.

## Author Contributions

A.D.-M. designed the study, performed most of the experiments, analyzed the results and wrote the manuscript. R.E. performed experiments on selected mutants, analyzed activity assays and NGS sequencing results and contributed to manuscript writing. G.M. performed preliminary experiments. R.S. analyzed the results and contributed to manuscript writing. Y.R. designed and supervised the study, analyzed data and wrote the manuscript. All the authors discussed the results and commented on the manuscript.

## Methods

DNA strands for the molecular program and probes were purchased from Biomers, primers from Eurofins and rhqPCR primers and genes from IDT. *Bst* 2.0 WarmStart®, *Taq* and *OneTaq® Hot Start* DNA polymerases as well as the control nicking enzymes *Nt.BstNBI* were purchased from New England Biolabs (NEB). Details on sample preparation, DNA sequences, experimental conditions, NGS analysis and simulations are provided in the Supplementary Information.

### Isothermal primers amplification and PCR

The IPA (Isothermal Primers Amplification) reactions were performed in IPA_v0_ buffer (Tris-HCl pH 8.8 20mM, (NH_4_)_2_SO_4_ 10 mM, KCl 10mM, MgSO_4_ 6 mM, NaCl 20 mM) with 200 μM of each dNTPs (NEB), 0.1% of *Bst* 2.0 WarmStart® DNA Polymerase (8,000 U/ml), and the molecular program templates at the following concentrations: sceT 2nM, aT 40 nM, ppT_F_ and ppT_R_ 20 nM. For assays with bacteria (100 bact/nL, OD_600nm_ =0.5), protected versions of the templates with biotin and three phosphorothioate backbone modifications on each extremity (3’ and 5’) were used, in addition to 6μg/mL of Streptavidin (NEB) and 0.125% of rLysozyme (Merck 27–33 kU/μl). The mixture was then incubated at 45°C for the indicated amount of time (generally between 15 and 45 min). When PCR was linked to the IPA step, 4% *OneTaq® Hot Start* DNA Polymerase (5,000 U/mL) was added to the reaction, as well as plasmids in the case of *in vitro* control (when no bacteria was present). Following the IPA step, an initial denaturation for 3 min at 95°C, followed by 20 PCR cycles were run with the following conditions: denaturation at 95°C for 10s, annealing at 65°C for 20 sec, extension at 68°C for 2 min. This PCR process was terminated with a final extension step for 5 min at 68°C.

### Library preparation

The library was created by error-prone PCR (Mutazyme II, Stratagene) following the manufacturer’s instructions with 50 ng of *dam* methylated NBI gene PCR product and 30 PCR cycles. The product was cloned into a pSB3K3 vector using either NEBuilder® HiFi DNA Assembly Cloning Kit (NEB) or the MEGAWHOP protocol^54^ based on a plasmid carrying the inactive variant E418A. Plasmid preparations of the libraries were transformed into home-made electrocompetent *E. coli* T7 express (NEB) containing a plasmid expressing the methylase Ple1. After 1h of recovery at 37°C, half of the transformed cells were grown overnight at 30°C in 5 ml LB broth containing ampicillin (100 μg ml^−1^) and kanamycin (30 μg ml^−1^), diluted 1:100 into 100 ml of fresh medium, induced with IPTG (50 μM) after reaching an OD_600nm_ of ~0.6 and incubated for another two hours at 30 °C. The culture was split into 1.5mL aliquots, centrifuged at 5,000 g for 5 min and the pellets were stored at −80°C.

### Selections

Frozen aliquots of induced bacteria were thawed on ice and washed twice by centrifugation (5 min, 5,000 g) with resuspension buffer (Tris-HCl pH 7.5, 50 mM, supplemented with 100 mM NaCl). The aliquots were finally resuspended in 100 μL of the resuspension buffer and the optical density of a 100-fold dilution was measured at 600 nm. A solution for the IPA-PCR was prepared as described above with addition of 0.4% w/v of pluronic f-127 and 1 ng/μL of yeast RNA for droplet interface coating. Half of the solution were mixed with bacteria (200 bact/nL) and the other half with rlysozyme (0.25% from stock). Droplets with a diameter of ~27 μm were produced (flow rates of 2 × 2μL min^−1^ for aqueous phases, pressure at ~500 mbar for the oil phase HFE 7500, 2% w/w EA surfactant, approx. 1 bacterium per droplet) at room temperature using a custom made 2 layer flow-focusing microfluidic device (inspired by ^55^). Collected droplets were then transferred in PCR tubes. In neutral conditions, the IPA step was run by incubating the droplets at 45°C for 30 min. For the kinetic stress (KS), the incubation time was reduced to 10 min while in the case of thermal stress conditions (TS) a pre-incubation step at 65°C for 3 min was added before the 30 min of IPA. The PCR was then launched using the program described above. Retrieved emulsions were broken using 0.5 emulsion volume of 1H,1H,2H,2H-perfluorooctanol. The selected pools were purified with Macherey and Nagel PCR clean-up kit and eluted with 15μL of ultra-pure water. 12μL of the products were then digested with 10U of DpnI in Cutsmart Buffer (NEB) in a 20μL reaction to remove the original plasmid. 2.5 μL of the digestion was PCR amplified in a 50μL reaction using 200 nM of internal primers (ipF and ipR) with the Q5® polymerase in Q5 reaction buffer with 200μM of dNTPs (NEB) with the following program : initial denaturation 98°C 30s, 25 cycles:98°C 5s, 65°C 15s, 72°C 1min, final extension 72°C 2min). After gel purification with Macherey and Nagel kit, the products were either re-cloned into the vector backbone and transformed into bacteria or sequenced using a MinION 1D^2^ FLOMIN107 flowcell, following manufacturer protocol. In the case of the mock selection experiment, the library was replaced by a mix of bacteria expressing either the wild-type enzyme or the inactive variant (E418A)^28^ with a 1:9 ratio. Selectivity was assessed by performing an rhqPCR (IDT) on the mock library before and after the selection.

