## Supplemental information for "Directed Evolution of Enzymes based on *in vitro* Programmable Self-Replication"

### Supplementary methods

- **Library transformation in electrocompetent cells**

T7 express® E. Coli cells were chosen for co-expression of the nicking enzyme and a cognate methylase (ple1) because they do not restrict methylated DNA (McrA-, McrBC-, EcoBr-m-, Mrr-). The bacteria were transformed with a high copy pUCIDT plasmid carrying the methylase ple1 under a leaky lac promoter following NEB protocol to protect genomic DNA NBI target site from nicking. The T7 express/ Ple1 cells were made electrocompetent. Briefly, a colony was grown overnight in 5 ml of LB-low salt medium with 100 µg/mL of ampicillin (37°C, 250 rpm) to reach an OD600 between 4 and 5. 2 mL of this culture were inoculated in 200 mL of LB-low salt medium with 100 µg/mL of ampicillin, the bacteria were then incubated at 37°C, 250 rpm until OD600 reached 0.6-0.8. The culture was rapidly chilled on ice for 15-20 min and swirled from time to time. It was then centrifuged at 8000 g for 5 min at 4°C in cold tubes and after removal of the supernatant the pellets were resuspended in 80mL of cold 10% glycerol solution. This step was repeated several times, reducing the volume of cold 10% glycerol used for the resuspension to 60 mL, then 40 mL and finally 2 mL. The final dilution was adjusted to obtain a final OD600 between 80 and 100 (0.8 to 1 for a 1:100 dilution). The cells were split in 200 µL aliquot into 1.5mL chilled tubes and stored at -80°C. To transform the libraries, 200 µL of home-made T7 express®/ ple1 electrocompetent cells were taken from the -80°C freezer and thawed on ice for 10 min. The DNA libraries were concentrated into 5 µL of ultra-pure water using Zymoresearch DNA clean & concentrator kit and added to the cells. The cells were transferred to a ice-chilled electroporation cuvette (2 mm large). The cuvette was then placed inside an electroporator (2500 V, 5 ms). 975 µL of SOC medium prewarmed at 37°C were added immediately before cells were incubated at 37°C for ~30 min in an orbital shaker. To assess the library size, dilutions were spread on LB Agar plates containing the proper antibiotic

(10 µg/mL of ampicillin and 30 µg/mL of kanamycin) and the plates were incubated at 30°C overnight.

- **Protein expression**.

500 µL of either the above transformed bacteria or a colony (for single clone protein expression) were diluted and grown overnight in 5mL LB medium with Ampicillin (100ug/ml) and Kanamycin (30 ug/ml), at 30°C and 250 rpm. The following day, cultures were diluted 1:100 and grown until OD600 reached 0.4-0.6. IPTG was added to final concentration of 0.4 mM and bacteria were incubated for 2 hours at 30°C and 250 rpm.

- **Protein purification**

Protein purification was performed using His SpinTrap from GE Helthcare. We used 50 mL of bacteria culture with the protein expressed. Bacteria were lysed via sonication on ice. After elution, we used Amicon Ultra 30K (Merck Millipore) to change buffer to 20 mM TrisHCl and 50 mM NaCl pH 7.4. Then diluted into same buffer with glycerol to a final 50% glycerol. We tested protein expression in a PAGE gel (NuPAGE 4-12% Bis Tris Gel Thermofisher) and quantified the yield by UV absorbance in a Clariostar.

- **Activity test**

To measure NBI activity we used a probe nicknamed *loopact* (See Fig. S9). It consists of a DNA strand forming a hairpin stem-loop. The double stranded stem displays the recognition site of NBI, and a quencher and fluorophore at the 3' and 5' extremities. When NBI cuts the top strand, it allows the separation of the quencher and fluorophore leading to an increase of the fluorescence which can be followed in real-time PCR machines. This probe has phophorothioate bonds at the 3' and 5' extremity, and in the short loop to avoid degradation in bacteria lysate. Activity tests using *loopact* were run in a solution of 1x Thermopol DF (prepared at the lab following New England Biolabs recipe), 20 mM NaCl, 2% BSA9000S, extra 6 mM $MgSO_4$, 250 nM of *loopact* probe and 120 bact/nl of cells with expressed NBI. The reaction is incubated at 45°C and fluorescence is measured every 30 sec.

- **rhqPCR**

The rhPCR was set up using DreamTaq buffer (1x), 1.5 mM of extra $MgSO_4$ (NEB), 200µM of dNTPs (NEB), 1% Evagreen (Biotium), 200nM of the two adapted rhPrimers (IDT DNA technologies, see table), 0.5% of DreamTaq DNA polymerase (Thermofisher) and 7.5 mU of

RNaseH2 (IDT DNA technologies). The 10 µL reactions were monitored in CFX96 Touch Real-Time PCR Machine (Bio-rad) upon 40 qPCR cycles (95°C 10 s, 52°C 20 s, 68°C 20 s) after 3 min of initiation at 95°C. Standards were run with plasmid of either active or inactive version of the gene starting from ~1 ng in 10 µL and diluted sequentially by 5 for 5 dilutions. To determine the proper amount of RNaseH2 giving the best dynamic range, various concentrations of RNaseH2 were tested for each reaction and each primer range. The results were compared to positive and negative control reactions performed with the corresponding classical PCR primers.

- **qPCR for Fig. S10.**

The qPCR was run in the same condition than the rhqPCR using the classical primers primF_qNBI and primR_qNBI (see table).

- **Inactive variant**

Site directed mutagenesis was used to generate the inactive version of the NBI gene E418A using the Q5® Site-Directed Mutagenesis Kit (NEB). The manufacturer protocol was used with the following primers: ATTTTTATTTGCAATGAAGCG and ACTGTATTTGCATGGCTTACGTG.

- **Molecular program monitoring**

For primer production monitoring assays 50 to 200 nM of probes were added to the IPA reaction. The 10µL reactions were monitored in CFX96 Touch Real-Time PCR Machine (Bio-rad) set up at 45°C and taking a measurement every ~30 s.

- **Magnesium optimization for molecular program (Fig. S2)**

The reactions were performed in IPA buffer (Tris-HCl pH 8.8 20 mM, $(NH_4)_2SO_4$ 10 mM, KCl 10 mM, $MgSO_4$ 5 mM, NaCl 20 mM ) with various added concentrations of $MgSO_4$ and 100 µM of each dNTPs (NEB), 1% of Nt.BstNBI (NEB 10000 U/mL), 1% of Bst DNA Polymerase Large Fragment (8,000 U/ml), the molecular program templates at the following concentrations: x11 10 nM, Cx11 40 nM, X11toON 40 nM. The mixture was then incubated at 45°C and fluorescence was monitored in CFX96 Touch Real-Time PCR Machine (Bio-rad).

- **PCR program modifications**

The PCR programs of the IPA-PCR were modified during the project. The first conditions were used during the development of the method and the second were optimized, after obtaining a functioning system, in order to challenge the reaction for primer concentration.

| Conditions | Init. Denaturation | Number of cycles | Denaturation | Annealing | Elongation |
|---|---|---|---|---|---|
| 1st | 95°C, 3min | 35 | 95°C, 10s | 56°C, 1min | 68°C, 20s |
| 2nd | 95°C, 3min | 20 | 95°C, 10s | 65°C, 20s | 68°C, 2 min |

- **Homology modeling and structural representation**

The structure of NBI have been modeled based on similarity with N.BspD6I (2EWF) sequence using Modeller9.11{Sali:1993ie}. Molecular graphics and analyses performed with UCSF ChimeraX, developed by the Resource for Biocomputing, Visualization, and Informatics at the University of California, San Francisco, with support from National Institutes of Health R01-GM129325 and the Office of Cyber Infrastructure and Computational Biology, National Institute of Allergy and Infectious Diseases{Pettersen:2021cm, Goddard:2018jf}. We performed a structural alignment of the NBI model with HindIII (2E52) to dock the DNA breaking site into NBI active site as in{Kachalova:2008hz}.

The aT template DNA model was generated with 3D-DART{vanDijk:2009ea} and docked in the NBI model by aligning NBI recognition domain with BpuJI recognition domain crystallized with its cognate DNA and Hind III restriction enzyme catalytic site (2E52) also crystallized with cognate DNA (as in same ref). The aT template was aligned to the cognate DNA scaffold of 2VLA at the recognition site level and the recognition site was then matched with the cleaving site of 2E52 cognate DNA.

- **MinION NGS sequencing**

We used kits EXP-PCB001 for barcoding, SQK-LSK108 for ligation, EXP-LLB001 for flowcell loading, and minION flowcell version FLO-MIN107, thus the sequencing was $1D^2$.

- **Selection Quality Index (SQI)**

The SQI formula for a selection process using one active and one fully inactive variant is : $SQIsel = \frac{\lambda e^{\lambda}(p'_{exp}-p)}{(e^{\lambda}-1)(1-p)}$ where λ is the mean number of bacteria per droplet, $p'_{exp}$ the proportion of active genes after one selection round and $p$ the proportion of active genes before the selection.

In the case of the mock library selection experiments (Fig. 4E), $\lambda = 0.4$ and starting with 10% of active genes ($p = 0.1$) we obtain *SQI* of 1.09, 1.05 and 0.89 for 15, 30 or 45 minutes of IPA incubation time, respectively.

- **NGS analysis**

We used Guppy 2.3.5 for basecalling (with dna_r9.4.1_450bps_flipflop.cfg configuration file) and demultiplexing. We kept only sequences containing one of the barcodes used experimentally. We used LAST{Kieibasa:2011do} for pairwise alignment to wild type reference sequence. We used a quality cut off parameter -e 5000 for the alignment. We chose a region to align longer than the ORF, to improve the detection of the extremes of the gene. Also, we used a curated set of reads (between 1700 and 1900 bases long and a mean Quality score >8) of the wild type to train LAST algorithm. Thus, the parameters for LAST alignment algorithm were:

| | A | C | G | T |
|---|---|---|---|---|
| A | 5 | -45 | -18 | -48 |
| C | -45 | 8 | -43 | -43 |
| G | -34 | -42 | 8 | -45 |
| T | -48 | -25 | -45 | 5 |

We used our own codes written in R and Mathematica for statistical analysis of sequences.

Nanopore sequencing has a high error rate (~5-10%), which makes harder the accurate analysis of mutagenic libraries containing only a few point mutations per gene. On the other hand, each read should be similar to a known sequence (the wildtype) with a few point mutations. This can be used to correct Nanopore sequencing data and improve the statistics on a gene library{Rocio:2020ea}.

Briefly, we first characterized the errors made by Nanopore sequencing in a set of reads of the wildtype. For each position and possible error, we fitted a classifier of correct/incorrect read as a function of the Qscore (Q) returned by the nanopore device. We then re-scaled the confidence value of each nucleotide read in the libraries as $p_{correct}(S) = p_{correct-nanopore}(S) \times p_{correct-fit}(S, P, n, Q)$, where $p_{correct-nanopore}(S) = 1 - 10^{-Q/10}$ for a read S, and $p_{correct-fit}(S, P, n, Q)$ is the value assigned by the logistic regression to read of nucleotide n on position P for the read S with Qscore Q.

DNA sequences were translated to amino acids using the genetic code, and the confidence values ($P_{correct}(A, p)$) were assigned as the product of the $p_{correct}(S)$ of the three bases on the codon.

We computed $f_L(A,p) = \sum_{seqs \in libraryL} p_{correct}(A,p) / \sum_{seqs \in libraryL, A \in aminoacids} p_{correct}(A,p)$, the normalized frequency of an amino acid A in a position p weighted by the confidence values, for library L.

The fitness effect of a mutation towards amino acid A, in library L, position p is then computed as $F_{eff} = log\big(f(L,A,p)/f(error - pronelibrary, A, p)\big)$. We restricted this calculation to mutations which appeared in at least 0.005% of the sequences of the library (weigthed by $P_{right}(A,p)$). The $F_{eff}$ values are provided in Supplementary Files as .xls documents.

### Supplementary table

| **Templates for IPA-PCR** | | |
|---|---|---|
| **Name** | | **Sequence (5’-3’) and modifications** |
| s | Signal oligonucleotide | CT**GAGTC**TTGG |
| aT | Autocatalytic template | CCAN**GACTC**NG-CCAN**GACTC**NG_Phos |
| aT *biot* | Protected autocatalytic template | biotin-*A*A*CCAN**GACTC**NG-CCAN**GACTC**N*G*A*-bioteg |
| sceT *biot* | Protected source template | biotin-*sp18-*C*C*AN**GACTC**NGGGTA**GACTC**TCGAG*A*T*A*T*CTCGA**GAGTC**TACC |
| ppT$_F$ *biot* | Protected primer-producing template Forward | biotin-*A*A*A*ATTCTGCCTCGTGATACGC-CCAN**GACTC**N*G*A*-biotin |
| ppT$_R$ *biot* | Protected primer-producing template Reverse | biotin-*A*A*T*TACTTCGCGTTATGCAGGC-CCAN**GACTC**N*G*A*-biotin |
| ppT_p17r3 *biot* | Protected primer-producing template for primer production measurement | biotin-*A*C*CATGGCACCACCACCA – CCAN**GACTC**N*G*A*-biotin |
| ppT_p25f2 *biot* | Protected primer-producing template for primer production measurement | biotin-*A*T*TAGCCATTAGATCACCTCCTTAGG – CCAN**GACTC**N*G*A*-biotin |
| x11 | Signal oligonucleotide | CA**GAGTC**CAAG |
| Cx11 | Autocatalytic template | C*T*T*GGACUCTG-CTTG**GACTC**TG_Phos |
| **Primers** | | |
| pF | Forward selection primer | GCGTATCACGAGGCAGAATTT |
| pR | Reverse selection primer | GCCTGCATAACGCGAAGTAAT |
| IpF | Internal primer forward | AGGATCCTAAGGAGGTGATCTAATG |
| IpR | Internal primer reverse | GATGGTGATGGTGGTCGAC |
| p25f2 | Forward primer | CCTAAGGAGGTGATCTAATGGCTAA |

| | | |
|---|---|---|
| | at the beginning of NBI gene | |
| p25r2 | Reverse primer in the middle of NBI gene (~700bp) | CAGATTTCCCTTCACGAATTTTACC |
| rhprimf_nbiON | Forward wild-type detection rhprimer | CCATTCCACGTAAGCCATrUCAAAC_3SpC3 |
| rhprimf_nbiOFF | Forward inactive variant detection rhprimer | CCATTCCACGTAAGCCATrGCAAAC_3SpC3 |
| rhprimr_nbi | Reverse rhprimer | ACAAGCAGATTTTAATGACCCrACGTCC_3SpC3 |
| primF_qNBI | Forward non selective primer | CCATTCCACGTAAGCCAT |
| primR_qNBI | Reverse non selective primer | ACAAGCAGATTTTAATGACCC |
| | **Reporting probes** | |
| Loopact | Protected activity reporting probe | CY5_A*T*C*TCAACT**GACTC**ACAGAC*T*T*T*T*T*G*<br>TCTGT**GAGTC**AGTTGAG*A*T*_BHQ2 |
| LoopactNuc | Protected non-specific activity reporting probe | FAM_*A*T*CTC AACTGAATCACAGAC*T*T*T*T*T*GTCT<br>GT**GATTC**AGTTGAG*A*T*_BHQ1 |
| p17 reporting probe | Double stranded reporting probe | CY5_TCGCATCCATGGCACCACC*A*C*C*A*A*Phos<br>+<br>T*G*C*CATGGATGCGA_BMN-Q650 |
| p25 reporting probe | Molecular beacon reporting probe | ROX_C*C*C*TAAACTTTAGCCATTAGATCACCTCCTTAGGTTTAGGG_BHQ2 |
| X11toON | Reporting template | T*T*A*CTCAGCCAAGAC-CTTG**GACTC**TG-TAMRA |

**Supplementary table 1. List of oligonucleotides used in this study.** The oligonucleotides containing the term '*biot'* in their name are protected against degradation in presence of cell lysate. 'Bioteg' indicates the presence of a biotin coupled to a triethyleneglycol linker, '*' corresponds to phosphorothioate bonds, 'Phos' corresponds to a phosphate group added to the 3'-end, 'sp18' corresponds to an hexaethyleneglycol spacer. The dashes '-' inside the DNA sequence help visualizing v11 binding sites. NBI recognition site GAGTC and its reverse complement GACTC are highlighted in bold font (mutation in red). N (indicating a random mixture of A, C, G and T) has been used on both bases flanking the recognition site, to avoid introducing extra-site specificity bias during the selection.

### Supplementary Figures

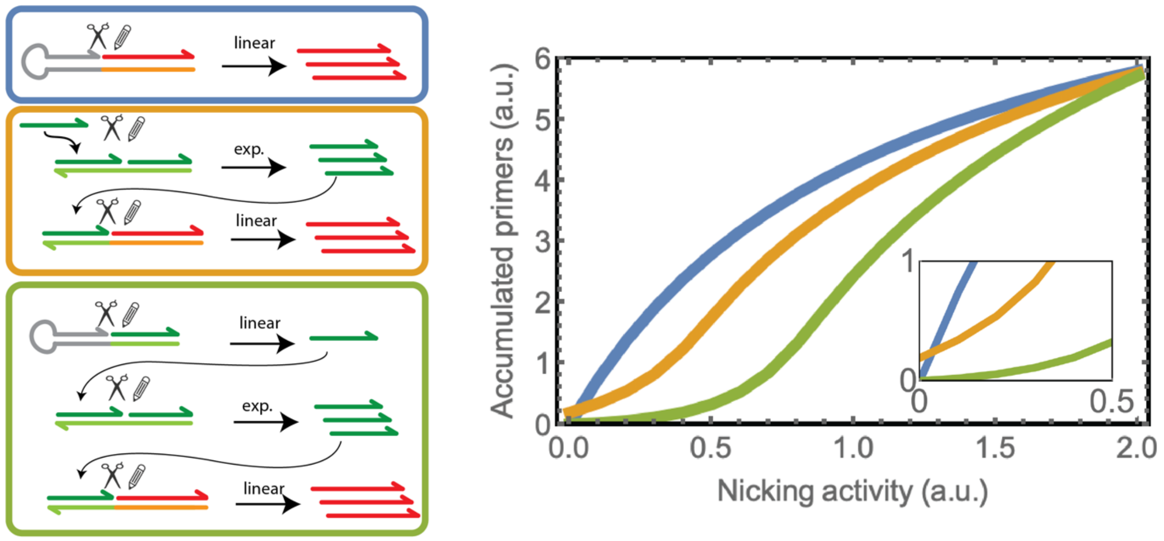


**Supplementary figure** 1**. Various possible designs for the selection circuits and simulation of the corresponding primer-response function.** The most basic design (blue frame and curve) directly uses NBI activity to release primers. However, this linear circuit has a tendency to accept variants displaying very weak activity, while providing diminishing returns to high activity variants. An additional exponential amplification stage can be inserted before primer production to obtain a sigmoidal response (orange frame and curve). But the presence of an initial small amount of trigger strands produces a baseline effect, enabling low level selection of inactive variants (inset). The third design (green frame and curve) avoids this issue by using a weak NBI-dependent production of the initial signal strands, which are then exponentially amplified and converted to primers. In this case, the network can efficiently reject low activity variants and provide an efficient (non-linear) response for the selection of active enzymes. The dynamic range can be tuned by playing on template concentrations or experimental conditions. Note also that the PCR stage is expected to bring additional non-linearity to the global selection (or fitness) function, since its response is also sigmoidal with respect to primer concentration.

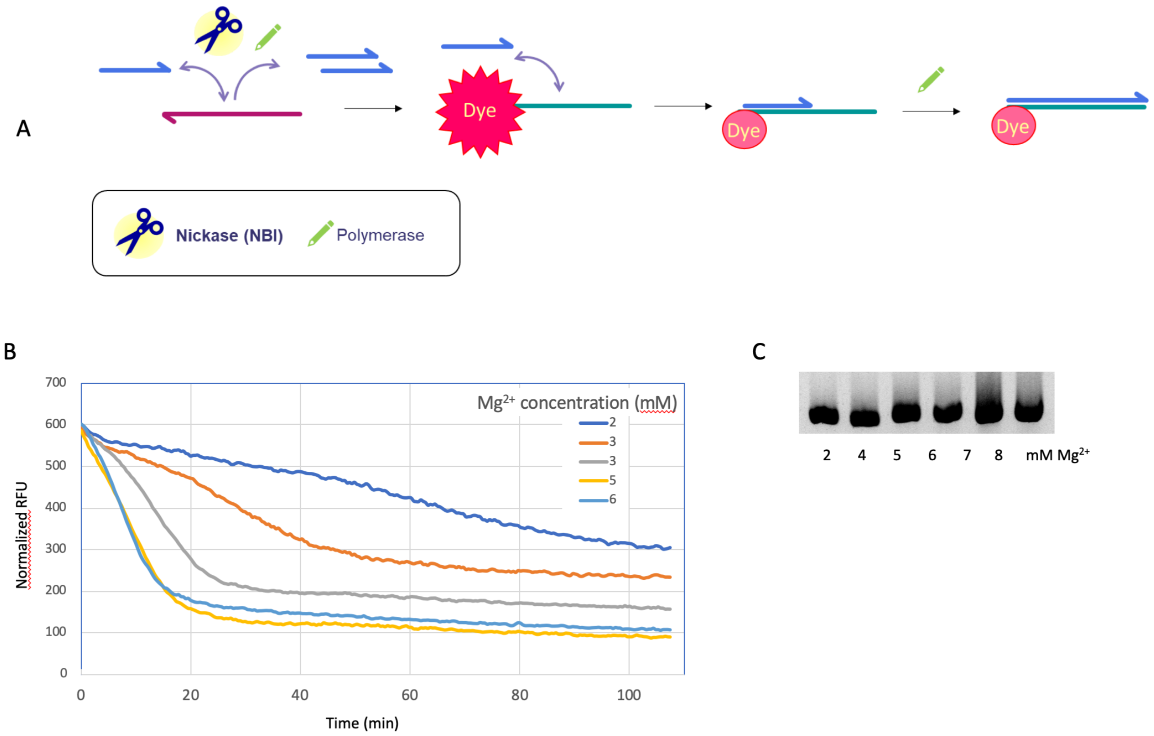


**Supplementary figure 2. Optimization of $Mg^{2+}$ concentration for compatibility of IPA and PCR reactions. A.** Scheme of the reporting system used. An autocatalytic strand (purple) binds an oligonucleotide signal (blue), which is exponentially replicated by a DNA polymerase and a nicking endonuclease. The signal accumulates and binds a reporting template, a DNA strand which includes the complementary sequence for the oligonucleotide and a fluorophore (TAMRA) at its 3' end. This duplex is then elongated by a DNA polymerase, yielding a stable duplex. The interaction between the fluorescent dye and the last DNA base on the opposite strand modifies the fluorescence properties, decreasing the signal{Padirac:2012bx}. **B.** Efficiency of IPA amplification according to $Mg^{2+}$ concentration. We used X11 as the input oligonucleotide, the autocatalytic template Cx11 and the reporting template X11toON (see DNA sequences in table S1). The curves correspond to various total $Mg^{2+}$ concentration in the Thermopol PCR buffer from NEB (which originally contains 2 mM $Mg^{2+}$). While a quick fluorescent decrease is observed at high $Mg^{2+}$ concentrations, the amplification kinetics deteriorates for lower $Mg^{2+}$. **C.** PCR ($1^{st}$ conditions) of our target gene with OneTaq polymerase (NEB) in Thermopol buffer supplemented with 20 mM NaCl, 2% BSA and increasing concentrations of magnesium salt. The polymerase OneTaq can perform PCR amplification of a gene-long amplicon in a buffer containing up to 8mM of $Mg^{2+}$.

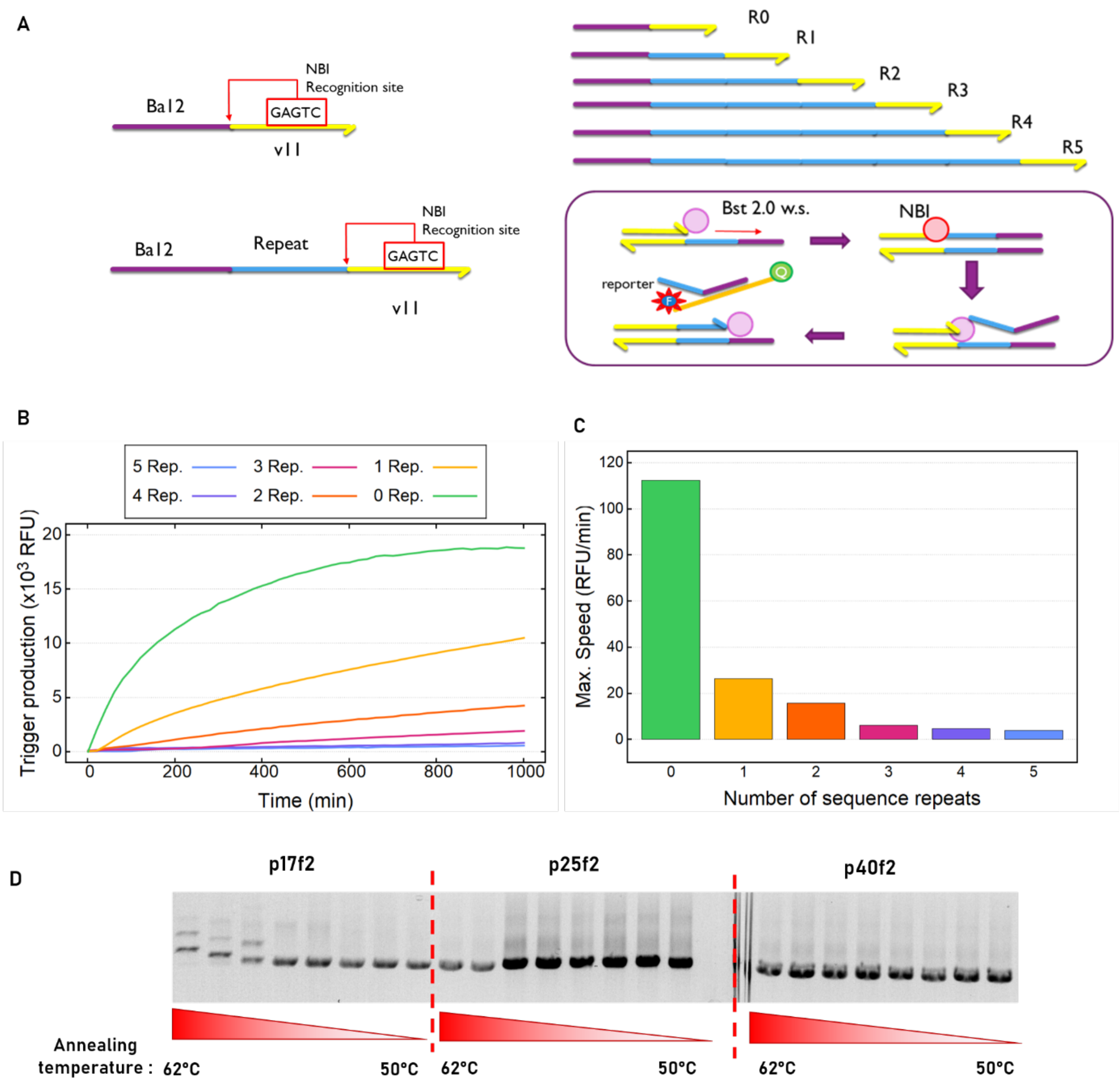


**Supplementary figure 3. Optimization of primer length for selection. A.** Description of the templates used for the experiment in panels B and C. 6 strands were designed containing a v11 binding domain (11 bases) sequence with NBI restriction site and Ba12 (12 bases) sequence that can bind specifically a random coiled single strand reporter (rBa12BQCy) containing on one side a quencher and a fluorophore (Cy5) on the other side. The difference between each strand is the number of repeats of a 10 bp sequence between Ba12 and V11. The experiment was run in 20 Mm Tris-HCl pH 8.8 ,10 mM $(NH_4)_2SO_4$, 10 mM KCl, 20 mM NaCl, 4.25 mM $MgSO_4$, 2% BSA9000, 3mM DTT, 0.025 % Bst WS 2.0, 2% NBI, 100 nM of v11 (s), 800 nM of rBa12BQCy reporter 100µM of dNTPs and 100 nM of v11toriBa12. **B.** Real time monitoring of the reporter fluorescence. **C.** Corresponding initial slopes. We observe that the rate of output production decreases quickly with the length of these outputs. **D.** PCR of NBI gene on pNBI plasmid with primers of various forward primer length (17 bp Tm = 46°C, 25 bp Tm = 59°C or 40 bp Tm = 64°C) and various annealing temperatures. The reverse primer is relatively short (17 bp) but with a 59°C Tm. Longer and more stable primers provide better amplification over a range of annealing temperature, although it seems still possible to use relatively short and weakly binding ones. Taking these two observations into account, we selected relatively short primers but with Tm around 58°C to perform the droplet selection. The final selection primers (pF and pR) had a Tm of ~58°C.

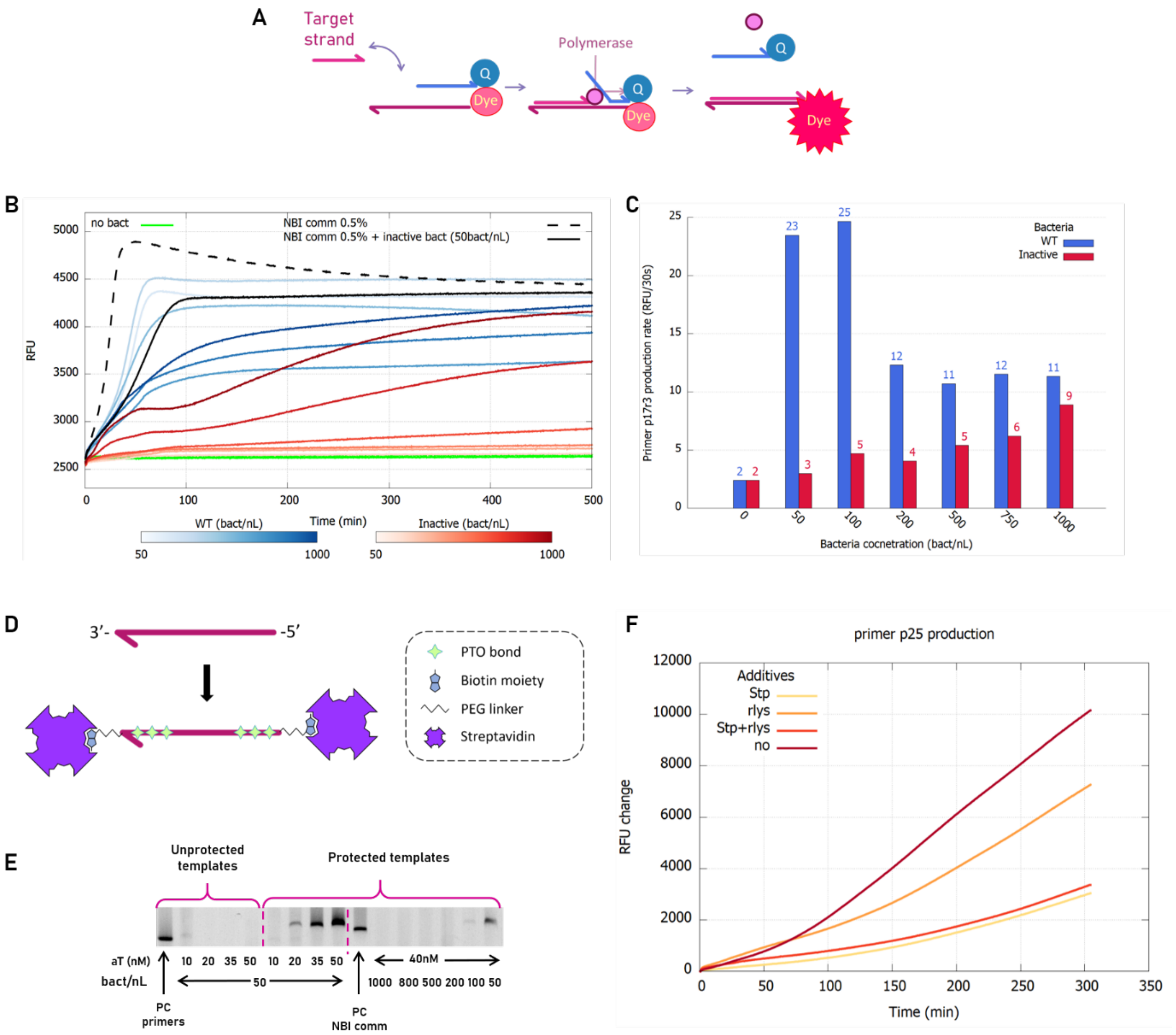


Supplementary figure 4. Templates protection. **A**. Scheme of the reporting system used to monitor primers production. The probe is constituted of a strand complementary to the target strand on its 3' extremity and carrying a 5' fluorophore, paired to a partial complementary strand carrying a 5' quencher. Upon binding and extension of the target strand, the quenching strand is displaced by the polymerase and the fluorescence is released. **B**. Real time monitoring of the production of primers with protected templates (s: 40 nM , aT : 40 nM , ppT_p17r3 biot : 20 nMm, ppT_p25f2 biot : 20 nM and p17 reporting probe : 250 nM) in the condition described for the IPA the presence of various concentration of lysate, from bacteria expressing either the wild-type NBI or the inactivated version. Purified NBI from NEB was used as a control (referred as NBI comm). **C**. Primer production rate as a function of bacteria concentration for bacteria expressing either the wild-type NBI or the inactivated version. **D**. Scheme representing the modification added to basic DNA oligonucleotides to protect them against degradation in presence of cell lysate {Tanno:2020ji}. **E**. Electrophoresis agarose gel obtained after IPA-PCR (1st conditions) for various autocatalytic template concentrations, various bacteria concentrations and using either unprotected or protected templates. **F**. Primer production along time using protected templates (s: 40 nM , aT : 40 nM , ppT_p17r3 biot : 20 nM, ppT_p25f2 : 20 nM and p25 reporting probe : 250 nM) in the condition described for the IPA in presence of rLysozyme (rlys) and/or Streptavidin (stp).

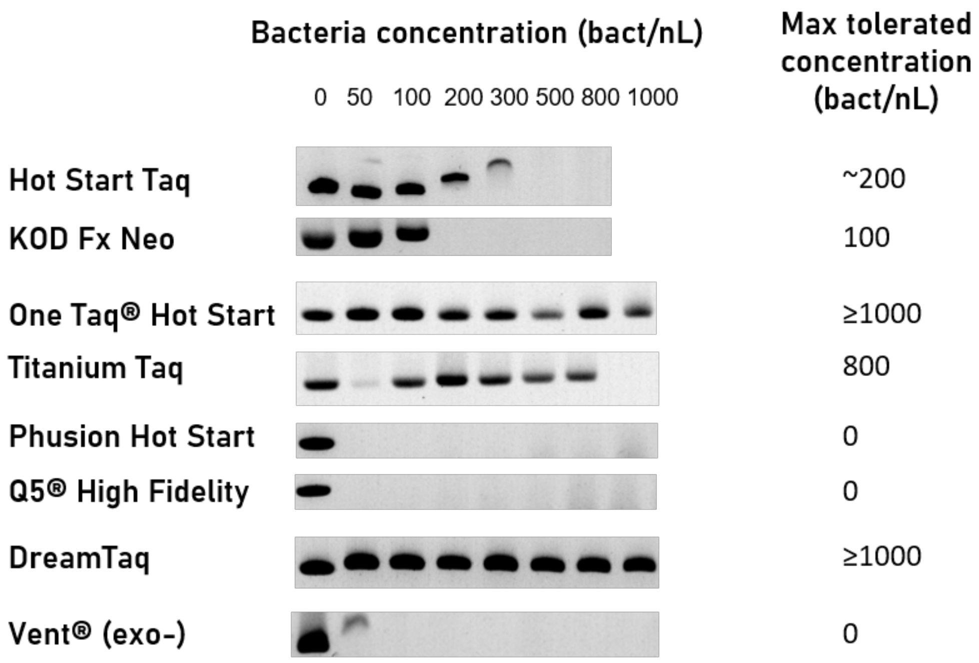


**Supplementary figure 5. Selection of a polymerase resistant to high bacteria lysate concentration.** We performed a PCR reaction with the primers p25r2 and p25f2, using a selection of commercially available DNA polymerases. We followed the protocol recommended by the manufacturer and used their respective commercial buffer and primer concentration, with increasing concentration of bacteria lysate. The initial DNA template was brought by the plasmids contained in the bacterial lysate (and thus its concentration was dependent of that of the lysate). For the 0 lysate controls, approximately 0.5 ng of miniprepped plasmid were spiked in each tube. DNA polymerases: Hot Start Taq (NEB), KOD Fx Neo (Toyobo), One Taq® Hot Start (NEB), Titanium Taq (Clontech), Phusion Hot Start (Thermofisher), Q5® High Fidelity (NEB), DreamTaq (Thermofisher), Vent (exo-) (NEB).

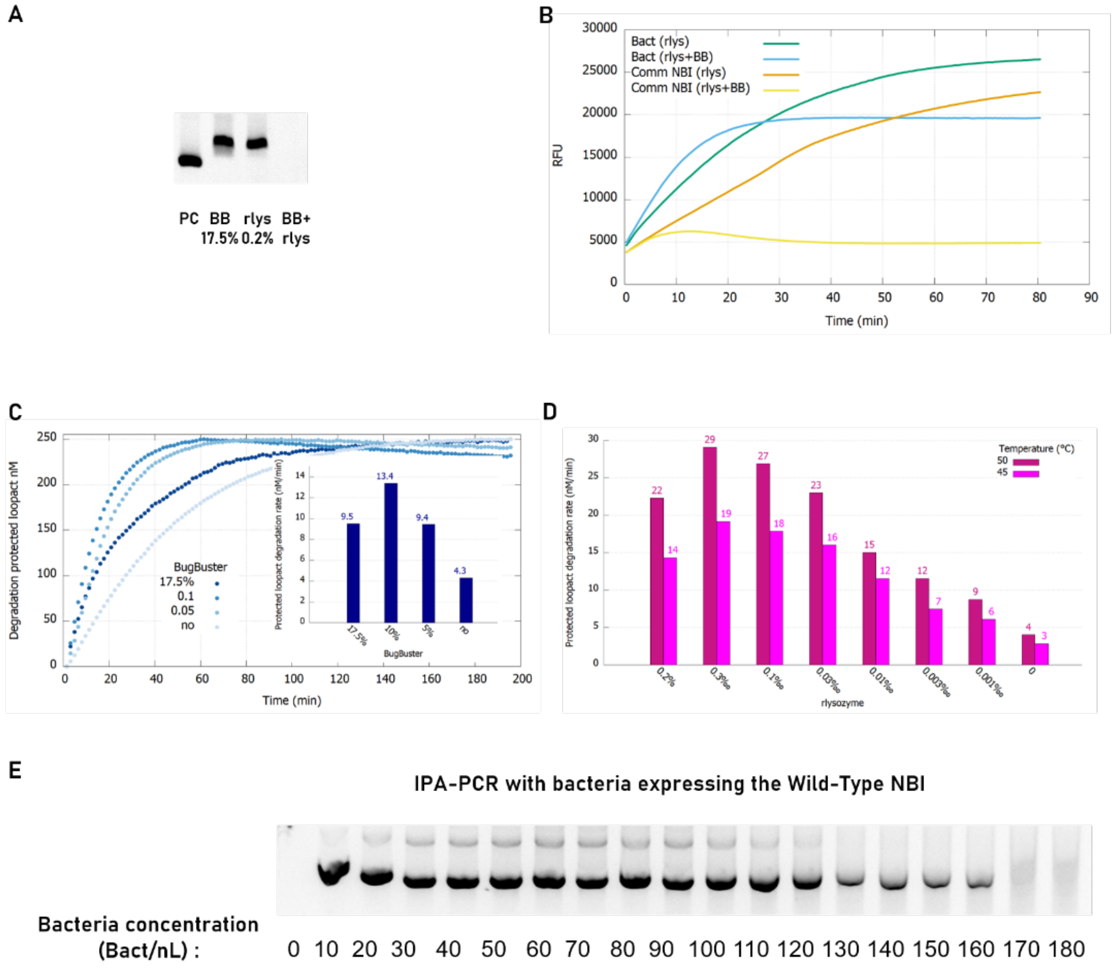


**Supplementary figure 6. Lysis conditions optimization for all-in-one selection process. A.** IPA-PCR reaction (1st conditions) run with the commercial NBI enzyme with 17.5% of BugBuster (BB) and/or 0.2% of rlysozyme (rlys). PC is the positive control without lysing agents. **B.** Activity assay of NBI either commercial (1%) or released from bacteria (100 bact/nL) using a the protected reporting probe *loopact* (cf. Supplementary Table 1) with 0.5% of rlys or 0.5% of rlys and 10% of BB. **C.** Effect of BugBuster concentration on NBI activity released by bacteria (50 bact/nL) in presence of 0.2% rlys at 45°C **D.** Effect of rlys concentration on NBI activity released by bacteria (50 bact/nL) at 45°C or 50°C, as measured by *loopact* digestion rate (see Fig. S9). **E.** IPA-PCR reaction (1st conditions) with 20 nM of trigger *s* instead of 2 nM of *sceT* template with increasing concentration of bacteria.

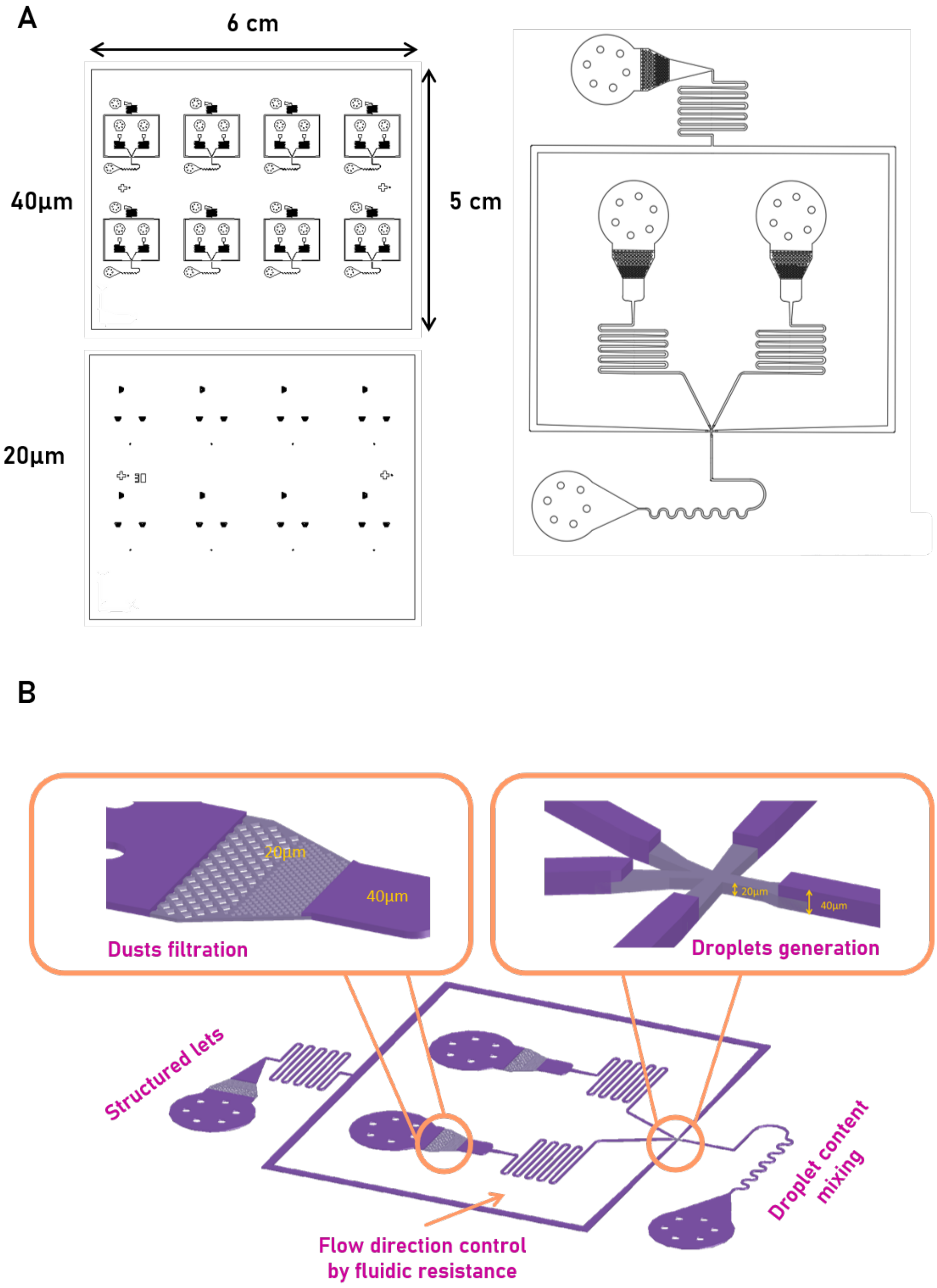


**Supplementary figure 7. Microfluidic device design. A**. A 2-step device with a first 20µm layer comprising the nozzle and the filters and a second 40 µm layer with the remaining part of the device. Marks are added for alignment. 8 devices can fit on a 50x60 mm PDMS slab that can be directly bounded to a microscopy glass slide of 72x56 mm. Pillars are added to the inlet and outlet punch area to avoid bonding of the PDMS ceiling on the glass. In the main part, channels are 50µm wide. **B**. Tridimensional representation of the final device.

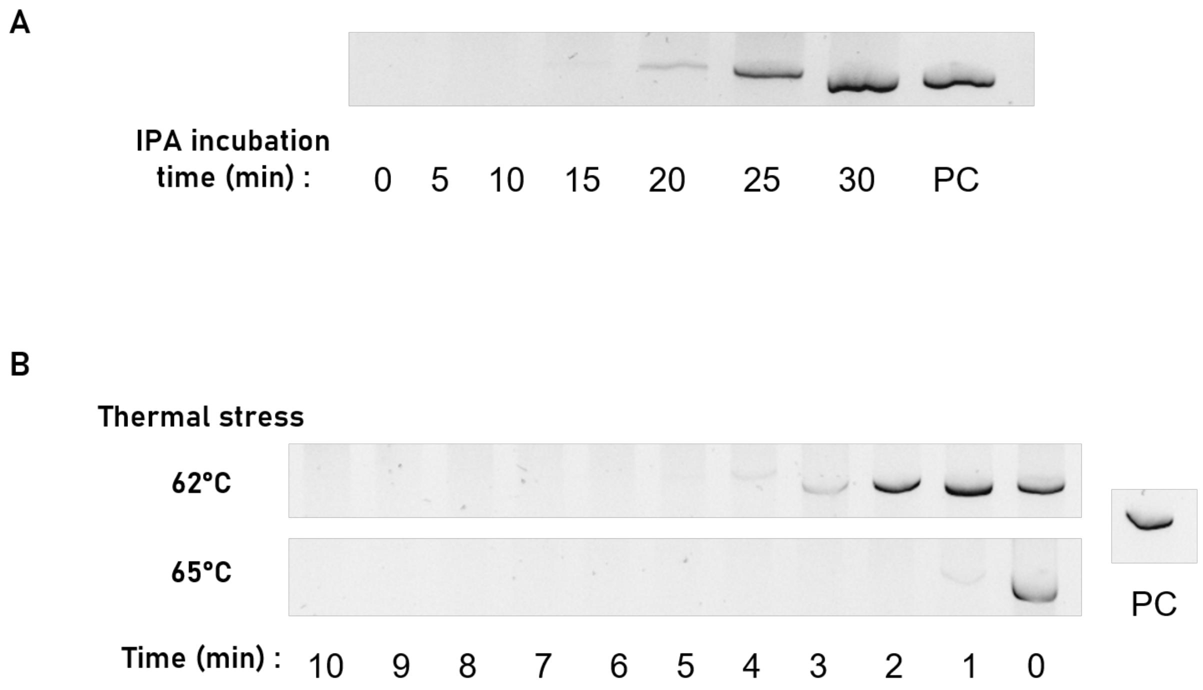


**Supplementary figure 8. Choice of the TS and KS selection conditions.** After optimization of the annealing temperature and duration to the 2nd and final conditions (cf Supplementary methods) in order to challenge the PCR reaction regarding the primer concentration we ran IPA-PCR reaction in order to determine the optimal conditions for selection. **A.** The IPA-PCR reaction was performed with various IPA incubation time. **B.** The IPA-PCR reaction was performed with a thermal stress of various durations either at 62 or 65°C, followed by a 30 min IPA incubation time.

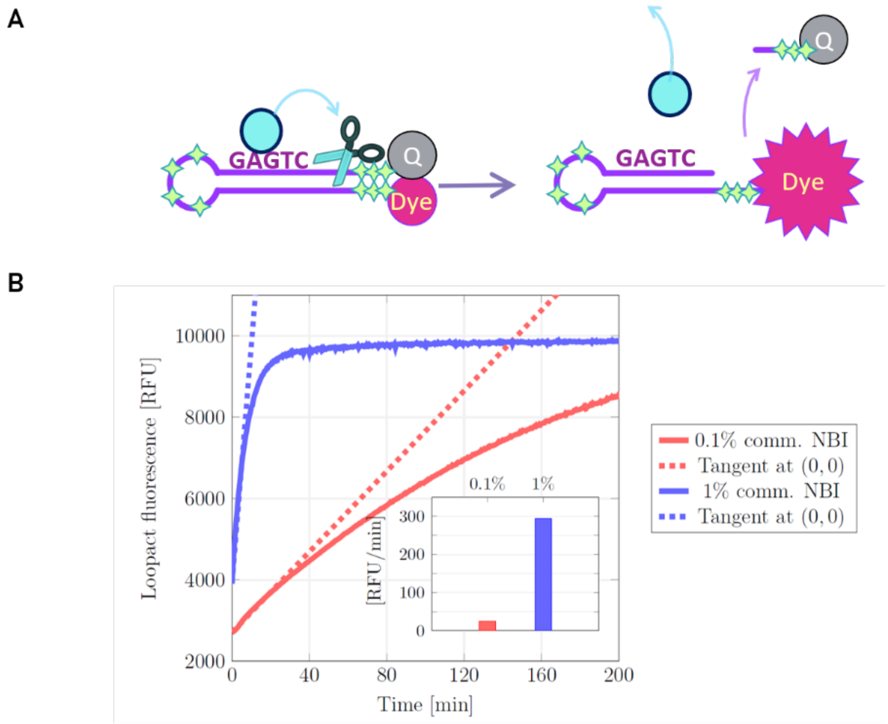


**Supplementary figure** 9**. NBI activity reporting assay. A.** Structure of the *loopact* probe, a DNA hairpin loop containing the site of the nickase and a fluorophore/quencher couple at the 5'- and 3'-ends. Once the strand is cut by NBI, the quencher melts away from the fluorophore, restoring fluorescence. Stars indicate phosphorothioate modifications for protection against non-specific degradation in the lysate. **B.** Real-time monitoring of *loopact* fluorescence in the presence of commercial NBI. **In inset,** the slope at origin for the two conditions.

A

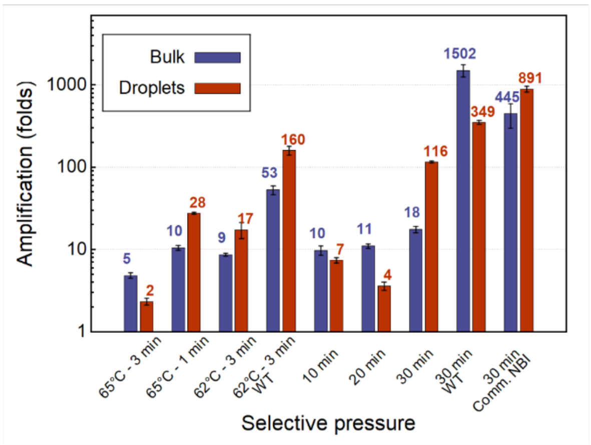


B

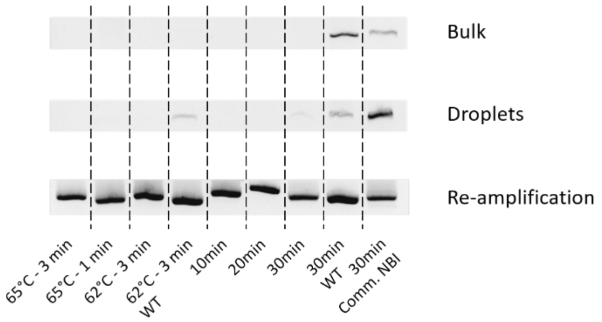


C

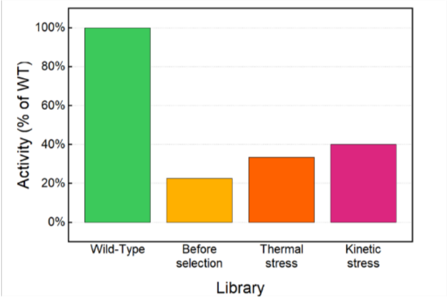


**Supplementary figure** 10**. qPCR Quantification and gel analysis of gene amplification during selection of the transformed library. A.** Gene amplification during IPA-PCR either in bulk (blue) or in emulsified (red) conditions for various selection protocols, as measured using the assay described in the supplementary methods. The first four conditions correspond to heat shocks at the given temperature for the given amount of time, prior to a 30 min-long IPA and subsequent PCR ($2^{nd}$ conditions). Condition described as TS in the main text is: 65°C – 3 min. The last five conditions correspond to various IPA durations. Condition described as KS and neutral in the main text are, respectively: 10 min and 30 minutes. 'WT' refers to the use of bacteria containing only the wild-type NBI gene, instead of the gene library. 'Comm. NBI' refers to the use of bacteria containing the inactive version of the NBI gene but in presence in the mixture of 0.3% of the commercial enzyme. Error bars corresponds to the standard deviation on duplicate measurements. In the case of the wild-type enzyme, or if the mixture is complemented with purified enzyme, we observe gene amplification (~3 orders of magnitude) in both droplet and bulk conditions. In the case of the library, since many bacteria contains inactive variants, the average activity is too low to support an efficient IPA-PCR process, and the bulk conditions show little amplification. For the droplet case, assuming that only droplets containing an IPA-active variant in one given condition should lead to an efficient droplet-PCR, the global amplification fold observed is expected to reflect, linearly, the fraction of active variants in that condition. **B.** Agarose electrophoresis gels after IPA-PCR for the selection conditions described before in bulk ('Bulk') and emulsified ('Droplets') conditions. The row 'Re-amplification' corresponds to the products obtained after PCR amplification on every 'Droplets' samples using nested internal primers (ipF and ipR at 200 nM, different from the ones used for the IPA-PCR). **C.** Measured lysate activity with the *loopact* protected probe for the library before and after the Thermal and Kinetic selections.

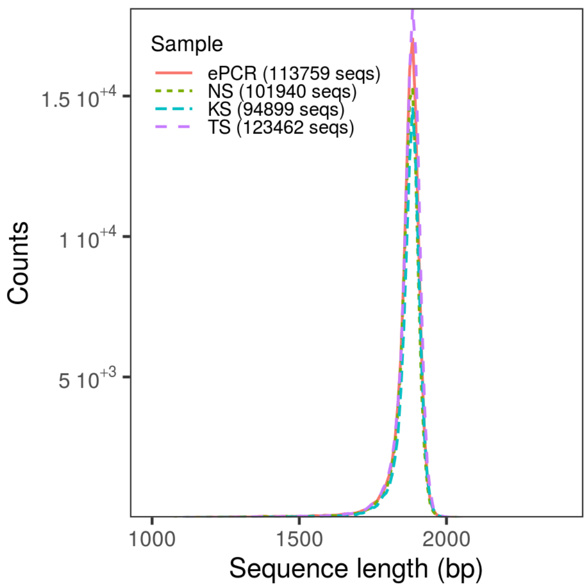


**Supplementary figure 11. MiniON reads statistics.** Length distribution of minION sequencing data, for ePCR (original mutagenic library), NS (subjected to neutral selection), KS (subjected to kinetic stress), and TS (subjected to thermal stress). Values computed after basecalling, demultiplexing and aligning to reference. For each set we obtained over $10^5$ sequences, and 434.060 sequences in total. The peak of the distribution is located at 1885 bases, close to the size of the reference sequence (which includes the ORF of the wildtype gene, plus the barcode used in nanopore sequencing protocol for multiplexing reads, 1927 bases).

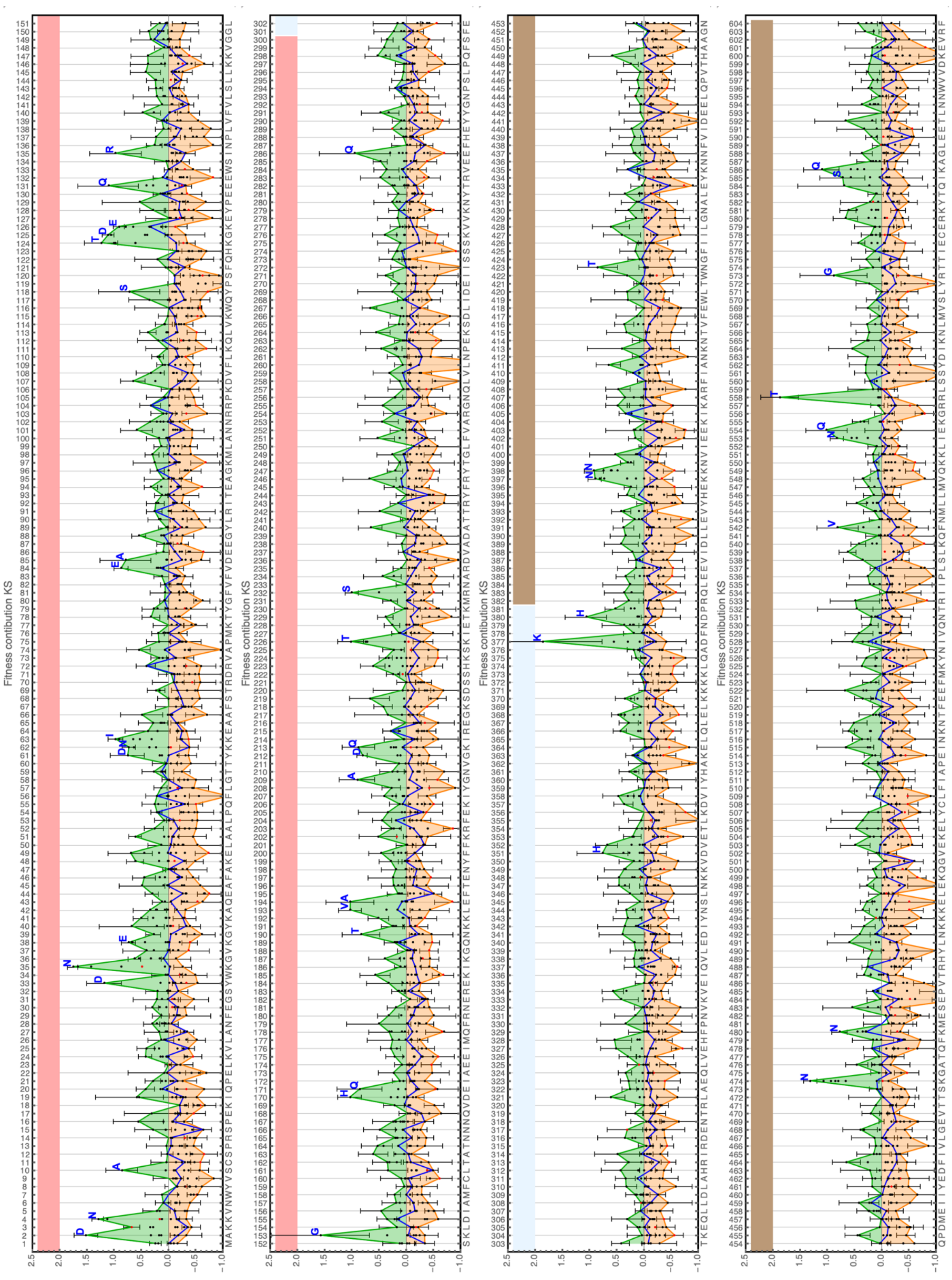


**Supplementary figure** 12. **Fitness contribution for the KS conditions.** All computed fitness contribution $F_{eff}$ are plotted as black dots along the protein sequence. For each position, the best and worst mutations are shown with their error bar, and the nature of the best mutation is indicated in blue when its score exceeds a threshold. All best mutations are connected by a green line, all worst mutations by an orange line and the neutral mutations by a blue line. Stop codon mutations are shown as red dots. The pink, blue and brown strips indicate the catalytic, linker and binding domain respectively.

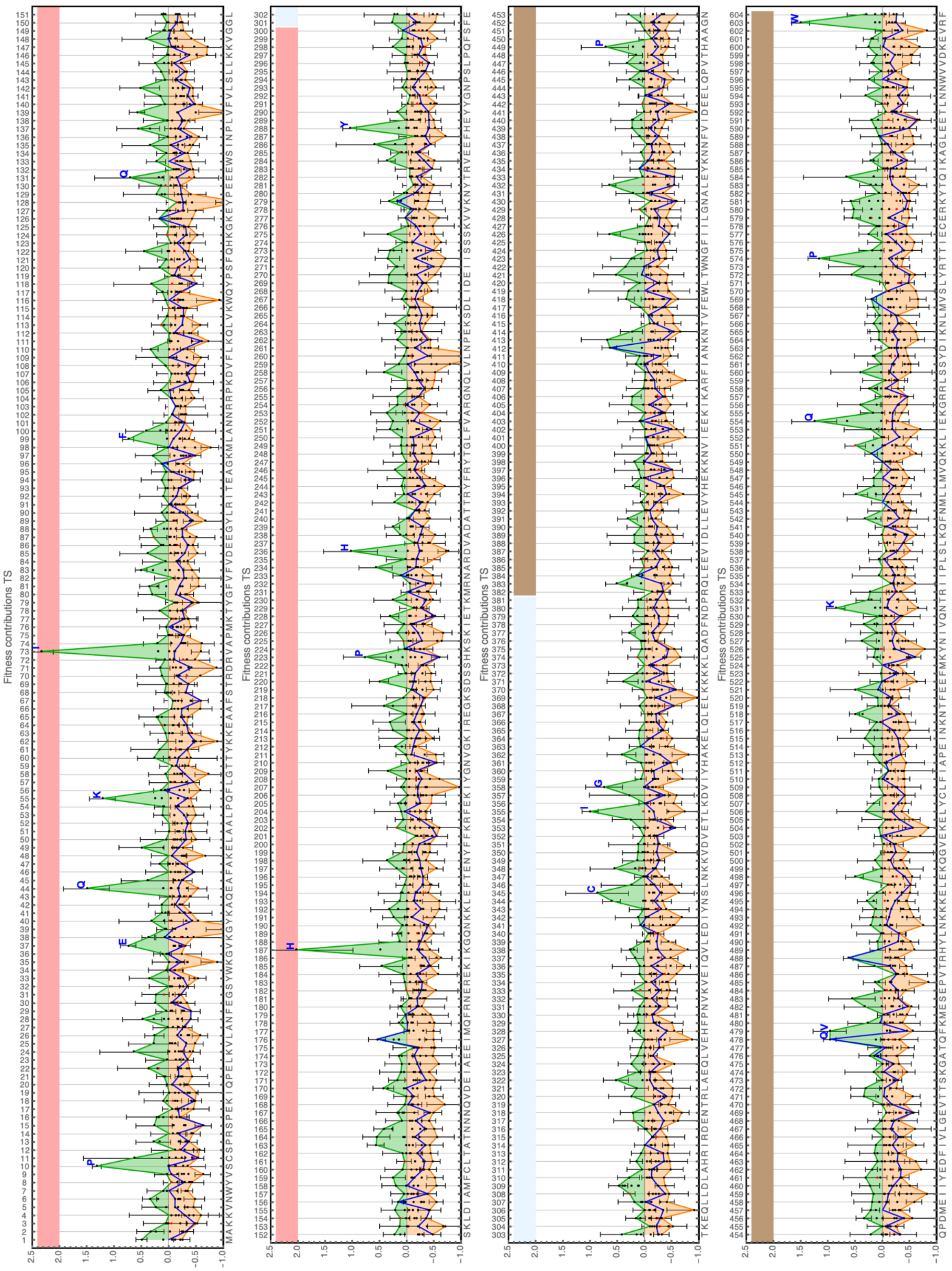

**Supplementary figure** 13**. Fitness contribution for the TS conditions.** Same as Fig S11 but for thermal shock conditions.

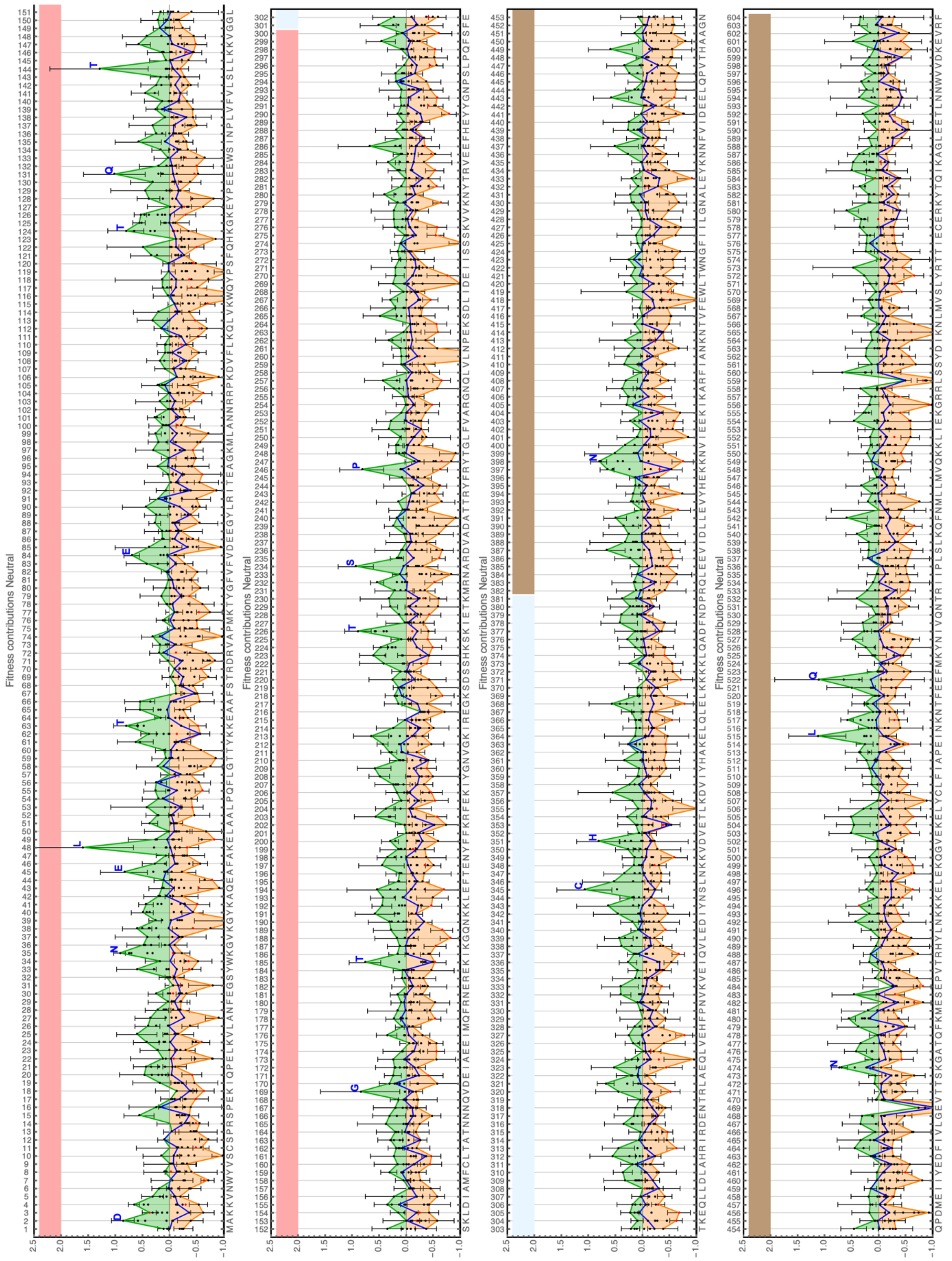


**Supplementary figure** 14**. Fitness contribution for the Neutral conditions.** Same as Fig S11 but for neutral conditions.

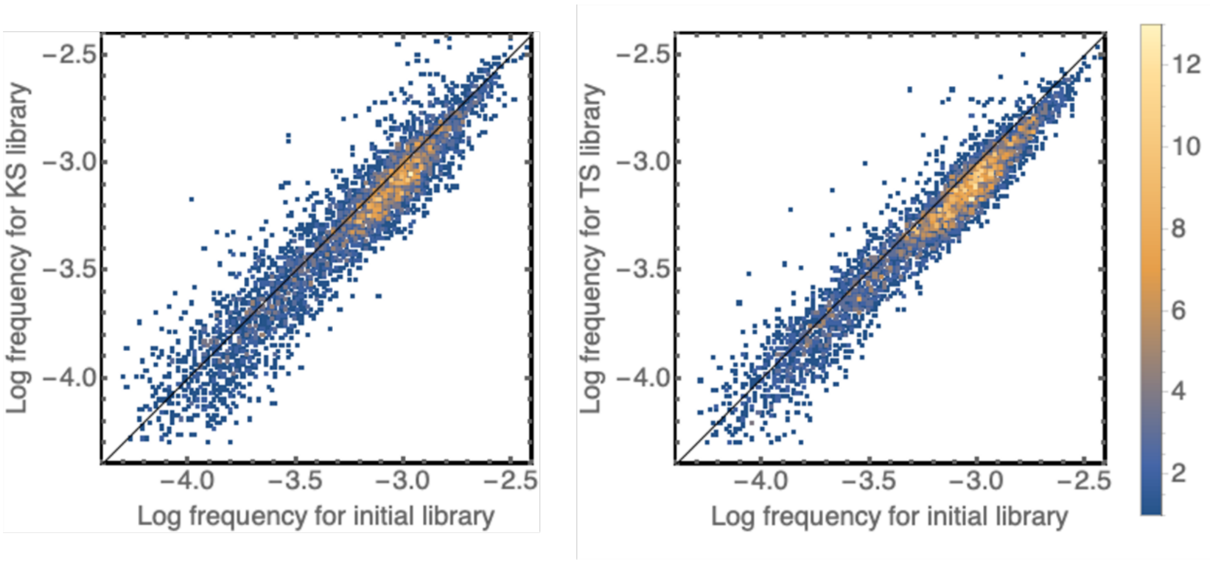


**Supplementary figure 15. Frequency change for the KS and TS selections.** The frequency of 3236 and 3368 mutations, respectively for the KS and TS protocols, in the initial and selected libraries. The frequency of mutations globally decreases during selections, but some of them show signs of positive selection.

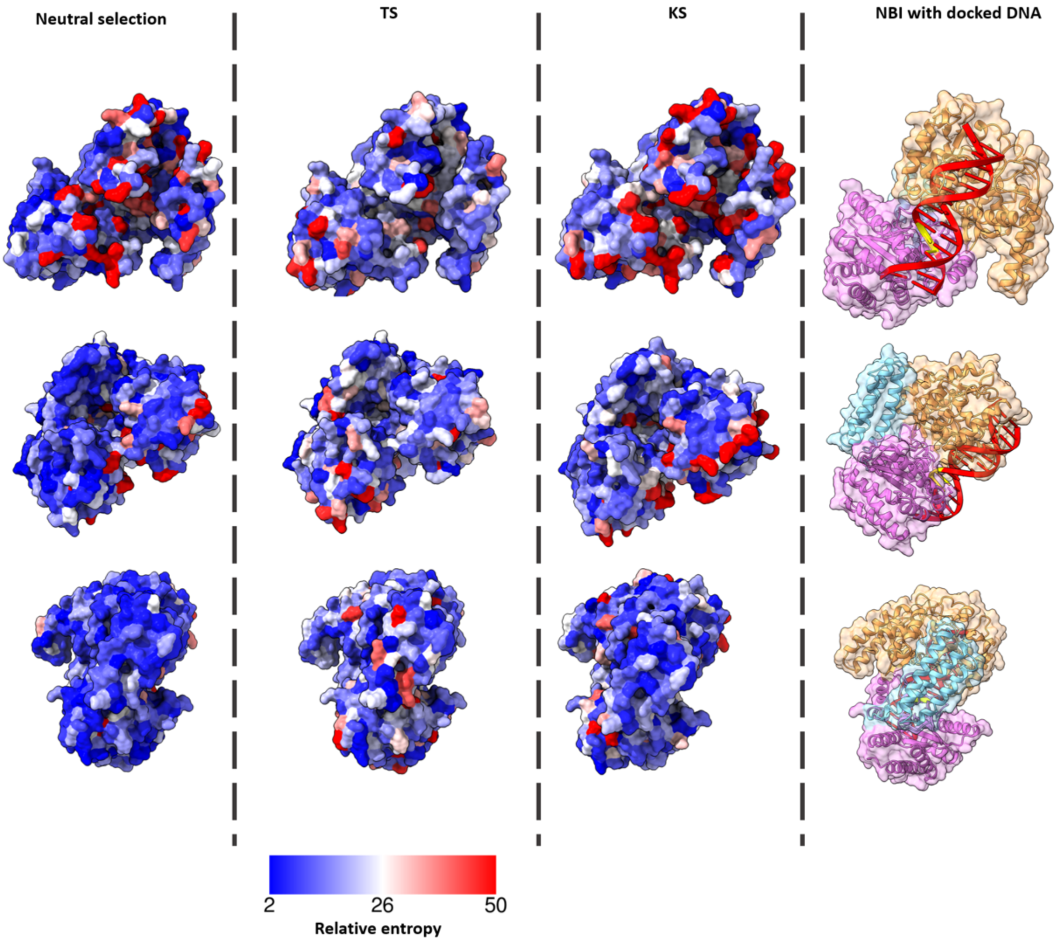


**Supplementary figure 16. Relative entropy of each position in different selection conditions.** Tolerance to mutations of each position was computed as the relative entropy of each position, as described in {Romero:2015bx}. Red represents strong response to mutations, blue weak response to mutations. On the right side NBI structure with docked aT template in the same orientation. In pink the catalytic domain, in blue the linker domain and in orange the recognition domain, the aT DNA template is in red with the cutting site in yellow.

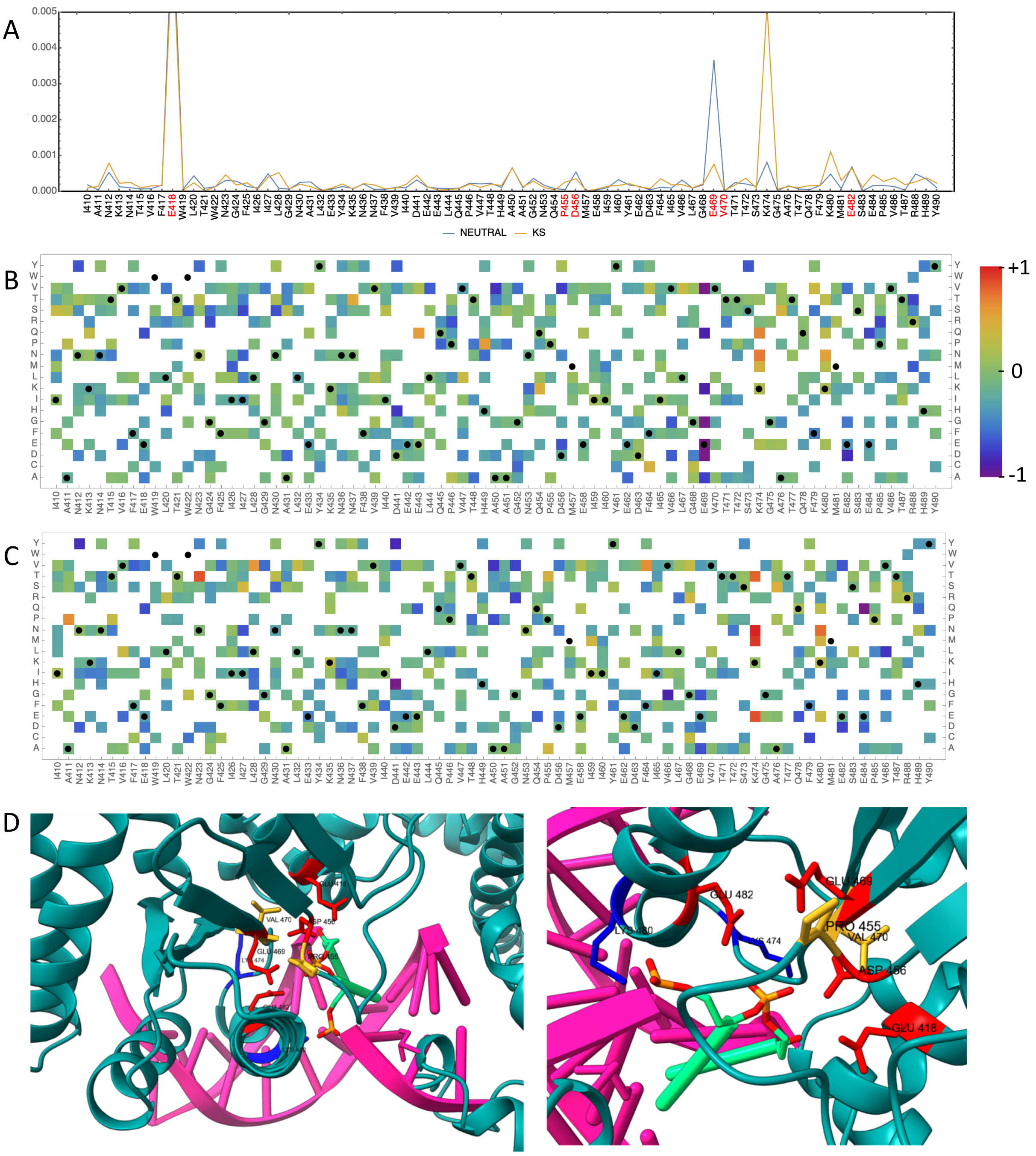


**Supplementary figure 17**. **Zoom on the active site residue for the neutral, and KS selections**. **A**. Relative entropy values. The putative active site residues are highlighted in red. **B**,**C**. Feff for the Neutral (left), and Kinetic selection (right), respectively. It has been reported that four negatively charged amino acids residues (E418, D456, E469 and E482) are essential for the catalytic activity of NBI. This stands in contrast to most type II restriction endonucleases, where the characteristic sequence of amino acid residues forming the catalytic center contains a positively charged lysine, following the motif PD...(D/E)XK, X being an hydrophobic residue{Kachalova:2008hz}. The sequence of NBI does display the characteristic PD (P455-D456), and EX (E469-V470) parts of this motif, but not the following lysine. Our scanning data (for the selections challenging the catalytic activity of the enzymes, i.e. the neutral and kinetic selections) provides support for the importance of the 4 negatively charged residues, but we do not observe strong effects associated with substitution of P455 or V470. In addition, we observe a strong selection effect on two nearby positively charged residues (K474 and K480). However, substitution of these amino acids by neutral of even negatively charged residues appears to provide beneficial effects. It is thus unlikely that they participate in the catalytic core, but may rather be involved in the general DNA-binding pattern (it was already noted that K474 does not structurally align with the active site lysin of similar restriction enzyme cleavage motifs e.g. FokI). Our results thus provide additional support for a fully negative catalytic site{Higgins:2001gl}, similar to the one of BamHI. Note that K474N is one of the substitutions we validated by directed mutagenesis, confirming its rate-accelerating effect (see main text Fig. 5), while E418A is the inactive variant that we have used. **D**. Zoom on a model of the active site of NBI (dark green) bound to DNA (pink). The NBI model active site domain was aligned to the HindIII restriction endonuclease (2e52) structure crystallized with the cognate DNA (RMSD 1.2 Å for aligned residues). This was used to visualize the position of the amino acids of the active site with respect to the cut site on DNA (light green). The reported essential negatively charged amino acids residues are in red (E418, D456, E469 and E482), the other residues of the characteristic motif PD...(D/E)XK are in yellow (P455 and V470), and the two rate accelerating positively charged residues are in blue (K474 and K480).

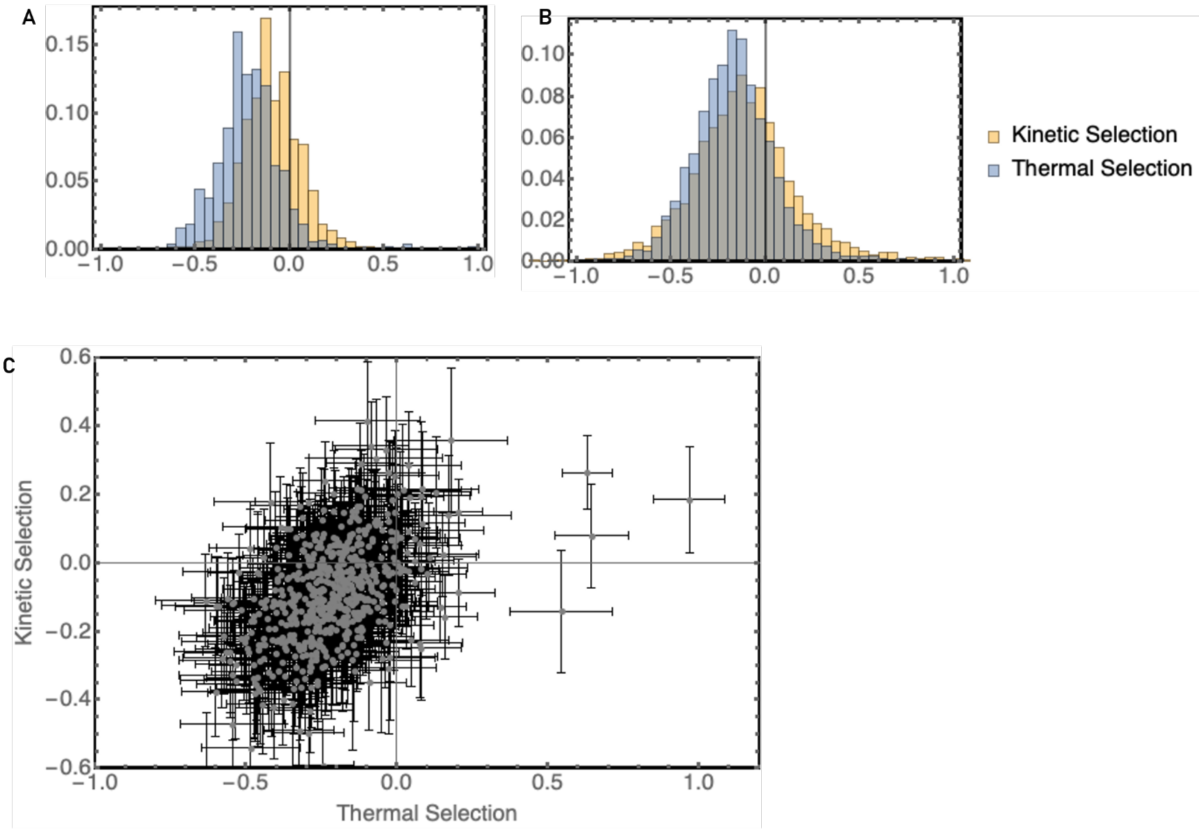


**Supplementary figure 18. Effect of synonymous mutations in KS and TS**: Distribution of fitness contributions $F_{eff}$ in the TS and Ks conditions. **A**: for synonymous mutation only. **B**: for all mutations. **C**. Scatter plot showing the effect of synonymous mutations in KS versus TS.

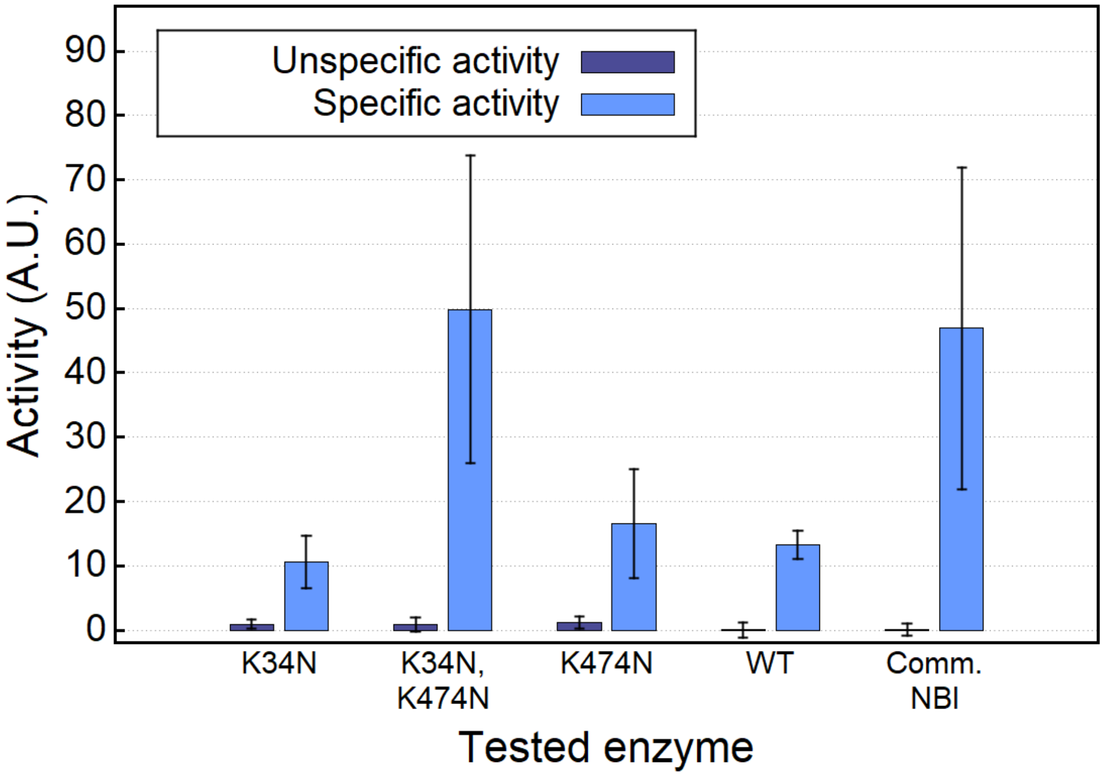


**Supplementary figure 19. Unspecific activity measurement of the mutated enzymes**. Enzymes activity measured with the specific *loopact* probe (containing the recognition site of NBI) and a nonspecific probe (*loopactNuc*, the same probe but with a mutation in the recognition site of NBI).

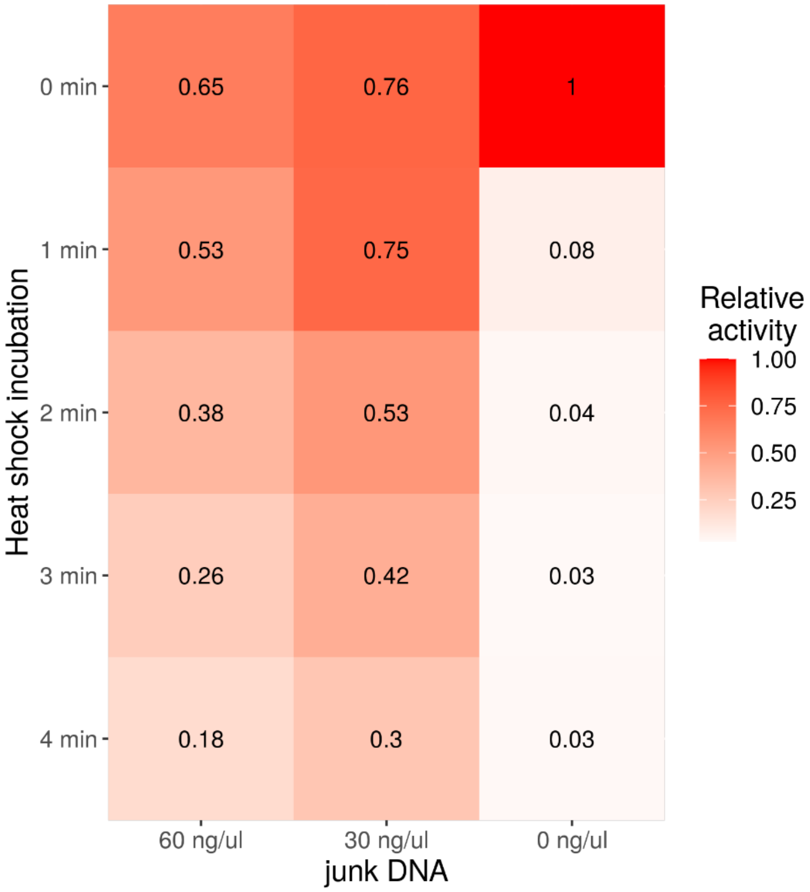


**Supplementary figure** 20**. Influence of extra nonspecific DNA on thermal resistance.** The activity of the commercial NBI enzyme assayed with the *loopact* probe (see Supplementary Fig. S9) with 0, 30 or 60 ng/µL of non-specific DNA (NBI gene PCR product, containing no NBI recognition site) after 0 to 4 min of 65°C heat shock. Non-specific DNA globally competes with the specific substrate, but also protects the enzyme from heat denaturation.

Ple1 plasmid sequence :

TCGCGCGTTTCGGTGATGACGGTGAAAACCTCTGACACATGCAGCTCCCGGAGACGGTCACAGCTTGTCTGTA
AGCGGATGCCGGGAGCAGACAAGCCCGTCAGGGCGCGTCAGCGGGTGTTGGCGGGTGTCGGGGCTGGCTTA
ACTATGCGGCATCAGAGCAGATTGTACTGAGAGTGCACCAAATGCGGTGTGAAATACCGCACAGATGCGTAA
GGAGAAAATACCGCATCAGGCGCCATTCGCCATTCAGGCTGCGCAACTGTTGGGAAGGGCGATCGGTGCGGG
CCTCATCGCTATTACGCCAGCTGGCGAAAGGGGGATGTGCTGCAAGGCGATTAAGTTGGGTAACGCCAGGGT
TTTCCCAGTCACGACGTTGTAAAACGACGGCCAGTGCAACGCGATGACGATGGATAGCGATTCATCGATGAGC
TGACCCGATCGCCGCCGCCGGAGGGTTGCGTTTGAGACGGGCGACAGATCCACAACATTTTGCGCACGGTTAT
GTGGACGAATTCGCGGCCGCTTCTAGAGTCGCGCGTTTCGGTGATGACGGTGAAAACCTCTGACACATGCAGC
TCCCGGAGACGGTCACAGCTTGTCTGTAAGCGGATGCCGGGAGCAGACAAGCCCGTCAGGGCGCGTCAGCGG
GTGTTGGCGGGTGTCGGGGCTGGCTTAAAGTCACGACGTTGTAAAACGACGGCCAGTGAAACCGAGCTCGGT
ACCCGGGGATCCTCAATAATTTGCAACAACTATATGCTTTGCAACAGACTTAAATCTATTCCTTATATTCACAGA
ATAACTTTTATCGTATTCATCAATAATCATTTTCTTATACAACCCATCAGTCAATTCAGTCTTACCGATAACCATC
ATGGCTTTGCATTTCAATTTCTTAAAGTCCTCACTCAATTTCACATGAGCATCTACATTGAAACCATCTTTATATT
CCTCATTACCATAATCACTAAATACGCAATCATAGGGAGGGTCTAGGAATATAAAATCATCTTTTCTCGCCATCT
TGAAGATCTCGCTGTAATCTTCATTAAATATCTGAGCACCCTGCATTAACAAGTGATGTTCATTAGCCACAAGTT
TTGTATTGAGATTTTTATATCTACCAAACGGAACATTAAACTCACCTTTTGAGTTATATCTAATCATTCCAGAGTA
CGCAGTCTTATTTATATAAAAATACAGTGTTGATGAAAGATATCTATCCGAAAAATCTTTATTAAATTCATTCCT
CATGAAGTAATAGAAATCTTCATTTCCATCATCTACACGTTCAGTAGGATTTAATTTCTTTCTAGTTTCGTATTCA
ACTCTATTCTTTTCATAAATACATTCTATCTCATCAAGCTCATGACGCAATTGAACAAAGTTATCTTTAACATCTC
GATAAAAATCTATAAGTTTTTTATTAATGTCATTGATAATAGATTTTTCTGGCTCTATATAAAAAAATAAAGCCC
CCCCACCAAAAAAAGGCTCTATGTAGCGCCCTTTAAATTCAGGGATATGTTTAATTAGATATGGAATTTCCTTA
GACTTTCCACCTCTATATTTAACTAATGGCTTCATACGATCACCTCCTTATCCGCAGGCATGCAAGCTTGGCGTA
ATCATGGTCATAGCTGTTTCCTGTGTGAAATTGTTATCCGCTCACAATTCCACACATTATACGAGCCGGAAGCAT
AAAGTGTAAACTAATGCAGGAGTCGCATAAGGGAGAGCGTCGAGATCCCGGACACCATCGAATGGCGCAAAA
CCTTTCGCGGTATGGCATGATAGCGCCCGGAAGAGAGTCAATTCAGGGTGGTGAATGTGAAACCAGTAACGT
TATACGATGTCGCAGAGTATGCCGGTGTCTCTTATCAGACCGTTTCCCGCGTGGTGAACCAGGCCAGCCACGTT
TCTGCGAAAACGCGGGAAAAAGTGGAAGCGGCGATGGCGGAGCTGAATTACATTCCCAACCGCGTGGCACAA
CAACTGGCGGGCAAACAGTCGTTGCTGATTGGCGTTGCCACCTCCAGTCTGGCCCTGCACGCGCCGTCGCAAA
TTGTCGCGGCGATTAAATCTCGCGCCGATCAACTGGGTGCCAGCGTGGTGGTGTCGATGGTAGAACGAAGCG
GCGTCGAAGCCTGTAAAGCGGCGGTGCACAATCTTCTCGCGCAACGCGTCAGTGGGCTGATCATTAACTATCC
GCTGGATGACCAGGATGCCATTGCTGTGGAAGCTGCCTGCACTAATGTTCCGGCGTTATTTCTTGATGTCTCTG
ACCAGACACCCATCAACAGTATTATTTTCTCCCATGAAGACGGTACGCGACTGGGCGTGGAGCATCTGGTCGC
ATTGGGTCACCAGCAAATCGCGCTGTTAGCGGGCCCATTAAGTTCTGTCTCGGCGCGTCTGCGTCTGGCTGGC
TGGCATAAATATCTCACTCGCAATCAAATTCAGCCGATAGCGGAACGGGAAGGCGACTGGAGTGCCATGTCCG
GTTTTCAACAAACCATGCAAATGCTGAATGAGGGCATCGTTCCCACTGCGATGCTGGTTGCCAACGATCAGAT
GGCGCTGGGCGCAATGCGCGCCATTACCGAGTCCGGGCTGCGCGTTGGTGCGGATATCTCGGTAGTGGGATA
CGACGATACCGAAGACAGCTCATGTTATATCCCGCCGTTAACCACCATCAAACAGGATTTTCGCCTGCTGGGGC
AAACCAGCGTGGACCGCTTGCTGCAACTCTCTCAGGGCCAGGCGGTGAAGGGCAATCAGCTGTTGCCCGTCTC
ACTGGTGAAAAGAAAAACCACCCTGGCGCCCAATACGCAAACCGCCTCTCCCCGCGCGTTGGCCGATTCATTA
ATGCAGCTGGCACGACAGGTTTCCCGACTGGAAAGCGGGCAGTGAGCGCAACGCAATTAATGTAAGTTAGCT
CACTCATTAGGCACCGGGATCTCGACCGATGCCCTTGAGAGCCTTCAACCCAGTCAGCTCCTTCCGGTGGGCGC
GGGGCATGACTATCGTCGCCGCACTTATGACTGTCTTCTTTATCATGCAACTCGTAGGACAGGTGCCGGCAGC
GCTCTTACTAGTAGCGGCCGCTGCAGGGGCCGTCGACCAATTCTCATGTTTGACAGATCAGTTCTGGACCAGC
GAGCTGTGCTGCGACTCGTGGCGTAATCATGGTCATAGCTGTTTCCTGTGTGAAATTGTTATCCGCTCACAATT
CCACACAACATACGAGCCGGAAGCATAAAGTGTAAAGCCTGGGGTGCCTAATGAGTGAGCTAACTCACATTAA
TTGCGTTGCGCTCACTGCCCGCTTTCCAGTCGGGAAACCTGTCGTGCCAGCTGCATTAATGAATCGGCCAACGC

GCGGGGAGAGGCGGTTTGCGTATTGGGCGCTCTTCCGCTTCCTCGCTCACTGACTCGCTGCGCTCGGTCGTTC
GGCTGCGGCGAGCGGTATCAGCTCACTCAAAGGCGGTAATACGGTTATCCACAGAATCAGGGGATAACGCAG
GAAAGAACATGTGAGCAAAAGGCCAGCAAAAGGCCAGGAACCGTAAAAAGGCCGCGTTGCTGGCGTTTTTCC
ATAGGCTCCGCCCCCCTGACGAGCATCACAAAAATCGACGCTCAAGTCAGAGGTGGCGAAACCCGACAGGAC
TATAAAGATACCAGGCGTTTCCCCCTGGAAGCTCCCTCGTGCGCTCTCCTGTTCCGACCCTGTCGCTTACCGGAT
ACCTGTCCGCCTTTCTCCCTTCGGGAAGCGTGGCGCTTTCTCATAGCTCACGCTGTAGGTATCTCAGTTCGGTGT
AGGTCGTTCGCTCCAAGCTGGGCTGTGTGCACGAACCCCCCGTTCAGCCCGACCGCTGCGCCTTATCCGGTAAC
TATCGTCTTGAGTCCAACCCGGTAAGACACGACTTATCGCCACTGGCAGCAGCCACTGGTAACAGGATTAGCA
GAGCGAGGTATGTAGGCGGTGCTACAGAGTTCTTGAAGTGGTGGCCTAACTACGGCTACACTAGAAGAACAG
TATTTGGTATCTGCGCTCTGCTGAAGCCAGTTACCTTCGGAAAAAGAGTTGGTAGCTCTTGATCCGGCAAACAA
ACCACCGCTGGTAGCGGTGGTTTTTTTGTTTGCAAGCAGCAGATTACGCGCAGAAAAAAAGGATCTCAAGAAG
ATCCTTTGATCTTTTCTACGGGGTCTGACGCTCAGTGGAACGAAAACTCACGTTAAGGGATTTTGGTCATGAGA
TTATCAAAAAGGATCTTCACCTAGATCCTTTTAAATTAAAAATGAAGTTTTAAATCAATCTAAAGTATATATGAG
TAAACTTGGTCTGACAGTTACCAATGCTTAATCAGTGAGGCACCTATCTCAGCGATCTGTCTATTTCGTTCATCC
ATAGTTGCCTGACTCCCCGTCGTGTAGATAACTACGATACGGGAGGGCTTACCATCTGGCCCCAGTGCTGCAAT
GATACCGCGAGACCCACGCTCACCGGCTCCAGATTTATCAGCAATAAACCAGCCAGCCGGAAGGGCCGAGCG
CAGAAGTGGTCCTGCAACTTTATCCGCCTCCATCCAGTCTATTAATTGTTGCCGGGAAGCTAGAGTAAGTAGTT
CGCCAGTTAATAGTTTGCGCAACGTTGTTGCCATTGCTACAGGCATCGTGGTGTCACGCTCGTCGTTTGGTATG
GCTTCATTCAGCTCCGGTTCCCAACGATCAAGGCGAGTTACATGATCCCCCATGTTGTGCAAAAAAGCGGTTAG
CTCCTTCGGTCCTCCGATCGTTGTCAGAAGTAAGTTGGCCGCAGTGTTATCACTCATGGTTATGGCAGCACTGC
ATAATTCTCTTACTGTCATGCCATCCGTAAGATGCTTTTCTGTGACTGGTGAGTACTCAACCAAGTCATTCTGAG
AATAGTGTATGCGGCGACCGAGTTGCTCTTGCCCGGCGTCAATACGGGATAATACCGCGCCACATAGCAGAAC
TTTAAAAGTGCTCATCATTGGAAAACGTTCTTCGGGGCGAAAACTCTCAAGGATCTTACCGCTGTTGAGATCCA
GTTCGATGTAACCCACTCGTGCACCCAACTGATCTTCAGCATCTTTTACTTTCACCAGCGTTTCTGGGTGAGCAA
AAACAGGAAGGCAAAATGCCGCAAAAAAGGGAATAAGGGCGACACGGAAATGTTGAATACTCATACTCTACC
TTTTTCAATATTATTGAAGCATTTATCAGGGTTATTGTCTCATGAGCGGATACATATTTGAATGTATTTAGAAAA
ATAAACAAATAGGGGTTCCGCGCACATTTCCCCGAAAAGTGCCACCTGACGTCTAAGAAACCATTATTATCATG
ACATTAACCTATAAAAATAGGCGTATCACGAGGCCCTTTCGTC

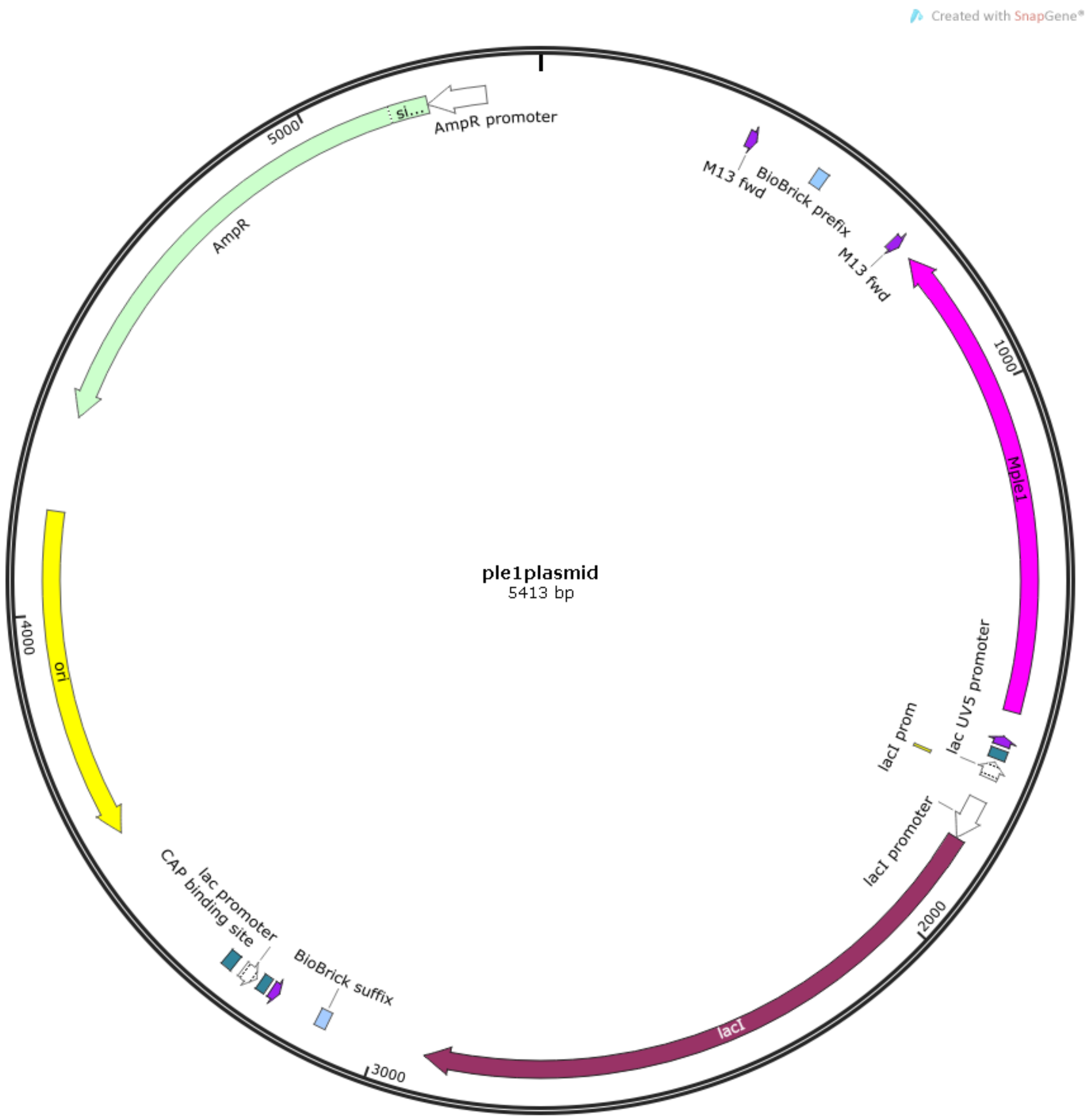
Created with SnapGene®
ple1plasmid
5413 bp
AmpR promoter
si...
AmpR
5000
M13 fwd
BioBrick prefix
M13 fwd
1000
Mple1
lac UV5 promoter
lacI prom
lacI promoter
2000
lacI
3000
BioBrick suffix
lac promoter
CAP binding site
ori
4000

pNBI plasmid sequence :
tgcaatttattcatatcaggattatcaataccatatttttgaaaaagccgtttctgtaatgaaggagaaaactcaccgaggcagttccataggatgg
caagatcctggtatcggtctgcgattccgactcgtccaacatcaatacaacctattaatttcccctcgtcaaaaataaggttatcaagtgagaaatc
accatgagtgacgactgaatccggtgagaatggcaaaagcttatgcatttctttccagacttgttcaacaggccagccattacgctcgtcatcaaa
atcactcgcatcaaccaaaccgttattcattcgtgattgcgcctgagcgagacgaaatacgcgatcgctgttaaaaggacaattacaaacaggaa
tcgaatgcaaccggcgcaggaacactgccagcgcatcaacaatattttcacctgaatcaggatattcttctaatacctggaatgctgttttcccggg
gatcgcagtggtgagtaaccatgcatcatcaggagtacggataaaatgcttgatggtcggaagaggcataaattccgtcagccagtttagtctga
ccatctcatctgtaacatcattggcaacgctacctttgccatgtttcagaaacaactctggcgcatcgggcttcccatacaatcgatagattgtcgc
acctgattgcccgacattatcgcgagcccatttatacccatataaatcagcatccatgttggaatttaatcgcggcctcgagcaagacgtttcccgt
tgaatatggctcataacaccccttgtattactgtttatgtaagcagacagttttattgttcatgatgatatatttttatcttgtgcaatgtaacatcaga
gattttgagacacaacgtggctttgttgaataaatcgaacttttgctgagttgaaggatcagatcacgcatcttcccgacaacgcagaccgttccgt
ggcaaagcaaaagttcaaaatcaccaactggtccacctacaacaaagctctcatcaaccgtggctccctcactttctggctggatgatggggcga
ttcaggcctggtatgagtcagcaacaccttcttcacgaggcagacctcagcgctagcggagtgtatactggcttactatgttggcactgatgaggg
tgtcagtgaagtgcttcatgtggcaggagaaaaaaggctgcaccggtgcgtcagcagaatatgtgatacaggatatattccgcttcctcgctcact
gactcgctacgctcggtcgttcgactgcggcgagcggaaatggcttacgaacggggcggagatttcctggaagatgccaggaagatacttaaca
gggaagtgagagggccgcggcaaagccgtttttccataggctccgcccccctgacaagcatcacgaaatctgacgctcaaatcagtggtggcga
aacccgacaggactataaagataccaggcgtttccccctggcggctccctcgtgcgctctcctgttcctgcctttcggtttaccggtgtcattccgctg
ttatggccgcgtttgtctcattccacgcctgacactcagttccgggtaggcagttcgctccaagctggactgtatgcacgaaccccccgttcagtcc
gaccgctgcgccttatccggtaactatcgtcttgagtccaacccggaaagacatgcaaaagcaccactggcagcagccactggtaattgatttag
aggagttagtcttgaagtcatgcgccggttaaggctaaactgaaaggacaagttttggtgactgcgctcctccaagccagttacctcggttcaaag
agttggtagctcagagaaccttcgaaaaaccgccctgcaaggcggttttttcgttttcagagcaagagattacgcgcagaccaaaacgatctcaa
gaagatcatcttattaaggggtctgacgctcagtggaacgaaaactcacgttaagggattttggtcatgagattatcaaaaaggatcttcacctag
atccttttaaattaaaaatgaagttttaaatcaatctaaagtatatatgagtaaacttggtctgacagttaccaatgcttaatcagtgaggcaccta
tctcagcgatctgtctatttcgttcatccatagttgcctgactccccgtcgtgtagataactacgatacgggagggcttaccatctggccccagtgct
gcaatgataccgcgagacccacgctcaccggctccagatttatcagcaataaaccagccagccggaagggccgagcgcagaagtggtcctgca
actttatccgcctccatccagtctattccatggtgccacctgacgtctaagaaaccattattatcatgacattaacctataaaaataggcgtatcac
gaggcagaatttcagataaaaaaaatccttagctttcgctaaggatgatttctggaattcgcggccgcttctagaTAATACAACTCACTATA
GGGGAATTGTGAGCGGATAACAATTCCCCTCTAGGAATAATTTTGTTTAGGATCCTAAGGAGGTGATCTAatgg
ctaaaaaagttaattggtatgtttcttgttcacctagaagtccagaaaaaaattcagcctgagttaaaagtactagcaaattttgagggaagttattg
gaaaggggtaaaagggtataaagcacaagaggcatttgctaaagaacttgctgctttaccacaattcttaggtactacttataaaaaagaagctg
cattttctactcgagacagagtggcaccaatgaaaacttatggtttcgtatttgtagatgaagaaggttatcttcgtataactgaagcagggaaaa
tgcttgcaaataaccgaagacccaaagatgttttcttaaaacagttagtaaagtggcaatatccatcgtttcaacacaaaggtaaggaatatccc
gaggaggaatggagtataaatcctcttgtatttgttcttagcttactaaaaaaggtaggcggcctcagtaaattagatattgctatgttctgtttaac
agcaacaaataataatcaggtggatgaaattgcagaggaaataatgcagttccgtaatgaacgtgaaaaaataaaaggacaaaataagaaac
ttgagtttactgagaattactttttaaaagattcgaaaagatttatggaaatgtaggtaaaattcgtgaagggaaatctgacagctcacataagt
caaaaattgaaactaaaatgagaaatgcacgagatgtggcagatgcaaccacaagatattttcgatatacaggtctatttgttgcaagagggaa
tcaactcgtcttaaatccagaaaaatctgatttaattgatgaaattatcagttcatcaaaagttgtaaagaactatacgagagtagaggaatttca
tgaatattatggaaatccgagtttaccacagttttcatttgagacaaaagagcaacttttagatctagcccatagaatacgagatgaaaataccag
actagctgagcaattagtagaacattttccaaatgttaaagttgaaatacaagtccttgaagacatttataattctcttaataaaaaagttgatgta
gaaacattaaaagatgttatttaccatgctaaggaattacagctagaactcaaaaagaaaaagttacaagcagattttaatgacccacgtcaact
tgaagaagtcattgaccttcttgaggtatatcatgagaaaaagaatgtgattgaagagaaaattaaagctcgcttcattgcaaataaaaatactg
tatttgaatggcttacgtggaatggcttcattattcttggaaatgctttagaatataaaaacaacttcgttattgatgaagagttacaaccagttact
catgccgcaggtaaccagcctgatatggaaattatatatgaagactttattgttcttggtgaagtaacaacttctaagggagcaacccagtttaag
atggaatcagaaccagtaacaaggcattatttaaacaagaaaaaagaattagaaaagcaaggagtagagaaagaactatattgtttattcattg
cgccagaaatcaataagaatacttttgaggagtttatgaaatacaatattgttcaaaacacaagaattatccctctctcattaaaacagtttaaca
tgctcctaatggtacagaagaaattaattgaaaaaggaagaaggttatcttcttatgatattaagaatctgatggtctcattatatcgaacaactat
agagtgtgaaagaaaatatactcaaattaaagctggtttagaagaaactttaaataattgggttgttgacaaggaggtaaggttttaagTCGAC
caccatcaccatcaccaccatcacccatggtaattaggtgattaaCTAGCATAACCCCTTGGGGCCTCTAAACGGGTCTTGAGG

GGTTTTTTGActagtagcggccgctgcagtccggcaaaaaaacgggcaaggtgtcaccaccctgcccttttctttaaaaccgaaaagattac
ttcgcgttatgcaggcttcctcgctcactgactcgctgcgctcggtcgttcggctgcggcgagcggtatcagctcactcaaaggcggtaatctcgag
tcccgtcaagtcagcgtaatgctctgccagtgttacaaccaattaaccaattctgattagaaaaactcatcgagcatcaaatgaaac

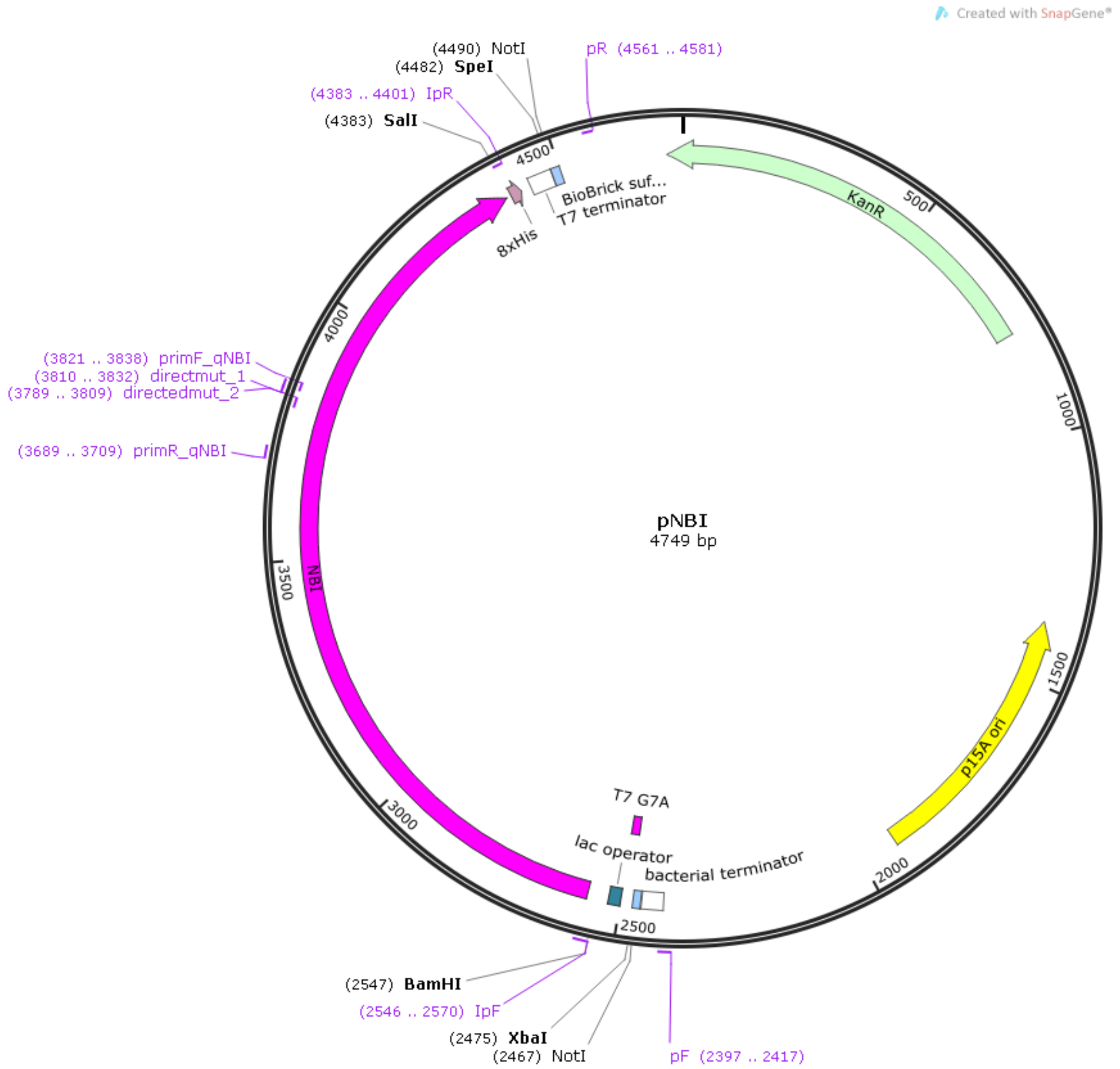
